# Neural integration of acoustic statistics enables detecting acoustic targets in noise

**DOI:** 10.1101/2025.05.29.656794

**Authors:** Artoghrul Alishbayli, Karol Przewrocki, Paul van Heumen, Bernhard Englitz

## Abstract

Sound detection amidst noise presents an important challenge in audition. Many naturally occurring sounds (rain, wind) can be described and predicted only statistically, so-called sound textures. Previous research has demonstrated the human ability to leverage this statistical predictability for sound recognition, but the neural mechanisms remain elusive.

We trained mice to detect vocalizations embedded in sound textures with different statistical predictability, while recording and optogenetically modulating the neural activity in the auditory cortex. Mice showed improved performance and neural encoding if they could sample the statistics longer per trial. Textures with more exploitable structure, specifically higher cross-frequency correlations improved behavioral performance as well as neural representation of background and vocalization. Activating parvalbumin-positive (PV) interneurons had an asymmetric effect, improving detection and neural encoding of vocalizations for low correlations, and impoverishing them for high cross-frequency correlations.

In summary, mice exploit stimulus statistics to improve sound detection in naturalistic background noise, reflected in behavioral performance and neural activity, relying on PV interneurons for temporal integration.

**Highlights:**

- Mice integrate statistical information indicated by behavior and neural activity
- Encoding of background sounds stays stable in A1, while vocalizations are enhanced
- High cross-frequency correlations improve target detection and neural encoding
- Activating PV cells improves detection of sounds with low cross-frequency correlations

## Introduction

The perceptual environment of animals is structured at multiple hierarchical levels and characterized by statistical regularities. Many theories of neural function postulate that the nervous system has evolved to exploit such regularities efficiently and the capability to reduce redundant information may optimize energy expenditure and survival (1–4). The suppression of redundant information usually manifests itself through the process of neural adaptation to constant, repetitive, or predictable stimuli (5). However, the details of circuit-level mechanisms that underlie this ability remain unclear.

In the auditory domain, one area where the functional role of adaptation has been prominent is in the context of a well-known psychological phenomenon known as the cocktail party effect (6,7): here, the signal-to-noise ratio is typically poor, but we are still able to attend to important information by filtering out irrelevant noise. This feature of the auditory system is thought to rest on its ability to adapt to sounds at multiple stages of processing, from cochlear hair cells (8) to the auditory cortex (9); for review see (10). Along these processing stages auditory responses exhibit progressively more noise invariance (11,12). Recent studies into the neural basis of such noise suppression have highlighted the auditory cortex as an important processing stage, where in particular GABAergic interneurons may play separate but complementary roles due to the differences in their connectivity and physiology (13–17).

Traditionally, adaptation to background noise has been studied using artificial stimuli such as white noise and pure tones that give researchers tighter experimental control (e.g. (14)). Such basic stimuli fail to shed light on features that are present in complex naturalistic stimuli. A promising set of sounds in this respect are sound textures (18). A sound texture is composed of a limited set of sound elements that overlap in time and frequency and exhibit time-invariant statistics and can be represented compactly in terms of their time-averaged statistical parameters (18). These parameters can then be used to synthesize textures that are perceived as similar to the originals by human listeners, suggesting that the statistics are sufficient to account for the perceptually relevant information (19). This raises the possibility that the auditory system may also represent such statistical regularities and use them to suppress the response to them (20,21) on the level of the auditory cortex. This could provide the neural basis for improved detection and recognition of embedded target sounds (22). However, most previous research in this domain has investigated sound textures in anesthetized or passive conditions, usually using non-parametric stimuli and outside the most relevant task context, i.e. an animal actively detecting target sounds amidst a challenging acoustic background (12,23–28).

Here, we study the neural encoding of conspecific target sounds in the context of a background sound texture including animal behavior and cell-type specific optogenetic modulation. We trained mice to detect mouse vocalizations embedded in sound textures with parametrically controlled statistics and recorded the responses of neural populations in the primary auditory cortex, which has previously been shown to be required for challenging target-in-noise detection (29). We hypothesized that if the auditory system had more time to sample the statistics of the sound textures, the detection and neural encoding of the target signal should improve. We find that mice can indeed detect vocalizations better if they had more time within a trial to sample the sound texture, consistent with a benefit of prolonged exposure to the statistical properties in this trial. Alongside, the neural encoding of vocalizations and their decodability improved with longer texture exposure inside a trial. The encoding of the acoustic textures transitioned from being based on the details of a realization to the sound statistics, matching with human behavior (30).

Next, we varied the correlation between frequency channels - a major source of redundancy in natural sounds - in the sound textures to check if the behavior and neural activity in response to the target sound is sensitive to this statistical property. Mice showed better performance and clearer neural representation of vocalizations in the context of high cross-frequency correlations (CFC) conditions, analogous to classical studies with comodulated maskers (CMR, (31,32). We find a double-dissociation of modulating PV interneuron activity: increasing PV activity improved both detection behavior and neural encoding for low CFCs contexts, while the opposite occurred in high CFC contexts. Activation of SST interneurons did not substantially change the behavior or neural encoding. The present results highlight the processing of complex natural stimuli in the auditory cortex and identify a specific role for PV interneurons in narrow-band processing of stimulus statistics, while suggesting that they are not constructively involved in wideband modulation processing, such as in CMR.

## Results

We trained mice (transgenic PV::ChR2, n=7; SST::ChR2, n=3) to detect vocalizations embedded in sound textures in order to understand the dynamics of adaptation to such acoustic contexts and how long-term statistics affect the processing of textural sounds in the auditory cortex. On each trial, a sequence of conspecific, chevron-type vocalizations (33,34) embedded in a background sound texture was embedded after a period of exposure to background noise alone (**Figure 1A**). A set of carefully designed acoustic textures was used, which differed parametrically in their marginal distribution in frequencies and cross-frequency correlations (see Methods & Supplementary Figure 1). These statistical properties could be varied largely independently (18). Texture stimuli were drawn randomly per trial and served as challenging background sounds for the vocalization detection task. Vocalizations were presented at pseudorandomized spectral locations, to prevent the animal from a selective listening approach. The duration of pre-exposure was randomly varied from 0 to 6 seconds to make vocalization onset unpredictable in any given trial. Licking during the vocalization presentation was considered a hit trial and rewarded with water, via a water spout positioned close to their mouth. Pupil diameter and neural activity from the primary auditory cortex (A1) were recorded concurrently. Peristimulus time histogram (PSTH), stimulus reconstruction and nonlinear decoding were used to understand the sensitivity of the system to statistical features of the background textures (**Figure 1B**). PV and SST interneurons expressing channelrhodopsin-2 (ChR2) were identified using opto-tagging and stimulated in a random subset of the trials to investigate their role in the encoding of textural statistics (**Figure 1C**). The main property of the background textures that was investigated in this study was cross-frequency correlations which was varied independently from the frequency marginal statistics in different trials (**Figure 1D-G)**. Below, details on statistical analysis are reported in the text and indicated in the figure panels, however, omitted from the captions to keep those at a reasonable length.

**Figure 1.**
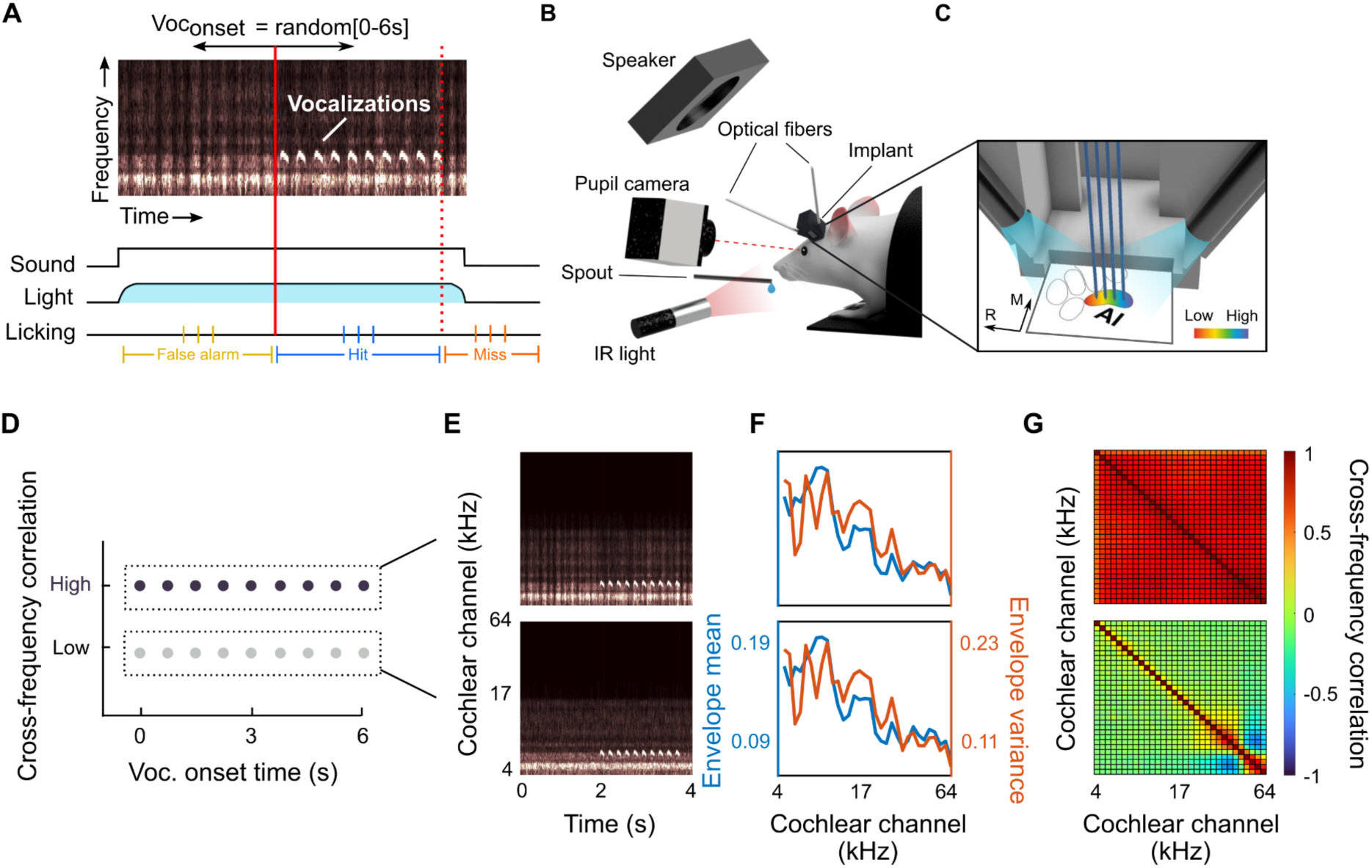
Neural recordings from the auditory cortex in mice actively detecting vocalizations masked by textural noise. **A** Mice were head-fixed during the training and neural recording sessions. This allowed them to be in the same position w.r.t. the speaker, placed centrally in front of the animal. During all sessions, the left eye was recorded while illuminated with infrared light. During neural recording sessions, neural activity was recorded from the left auditory cortex using microelectrode arrays with 64 Ch. and the brain surface could be illuminated with light (470 nm) to activate PV interneurons. **B** Electrodes were inserted at the primary auditory cortex such that they spanned along the tonotopic gradient in A1. The light was delivered bidirectionally through optical fibers that were inserted into the sides of the microdrive case. **C** In each trial the mouse listened for vocalizations embedded in a sound texture. The frequency and onset timing (0-6 s) of vocalizations were randomly assigned to different trials. In a subset of trials (30%, randomly selected) PV interneurons were slightly activated (∼5% change in FR) for the duration of the sound. The mice signaled that they detected a vocalization by licking, which led to a water reward if it occurred in the response window (0-1.9 s after vocalization onset), or no water if the lick occurred before (false alarm, FA), too late or not at all (miss). FAs were punished with a time-out of 7 s. **D** We tested the influence of cross-frequency correlations (CFCs) by using textures with low and high CFCs for multiple vocalization onset times. **E** Spectrograms of high and low cross-frequency correlation sounds. Sounds were matched for their marginal envelope mean and variance (**F**) but not for cross-frequency correlations (**G**). See Supplementary Figure 1 for all stimuli and Supplementary Figure 4C for all vocalizations.

### Mice learn to detect conspecific vocalizations embedded in textural noise

Head-fixed animals were trained for 4-6 weeks in the behavioral task, before being implanted with chronic electrodes. We adopted an individual training program for the mice to assist them in learning the task: multiple parameters of the paradigm were adapted to gradually make the task more difficult over time, e.g. by reducing the signal-to-noise ratio of vocalizations over background, extending the range of random onset times and shortening the response window. This training strategy prevented frustrating the animals early on and guided them to focus on the intended response strategy (detect vocalizations) over alternative response strategies (e.g. guessing or average-timing-based responses). We increased the difficulty such that the hit rate increased to and remained around 20-30%, thus keeping the task challenging for the mouse to motivate learning (35). To verify that each animal was responding to the vocalization, we estimated the chance performance by recomputing the chance hit rate in surrogate data, created by permuting the trial order (N=100) while keeping the original lick times and then taking the mean of the resulting distribution of chance hit rates (μ_rnd_). A session in which the animals performed significantly above chance, was defined as one where the actual hit rate was ≥ [μ_rnd_ + 2×σ_rnd_] (red dots in Fig. 2C; gray indicates insignificant). The performance ratio between actual and random performance served as an index for the learning progress (**Figure 2C**). Using a linear discriminant analysis (LDA), we found that the significant and insignificant performance ratios on the single recording level were best separated by a threshold performance ratio of >1.5. Using another LDA on the same data but separating along session number, we found that the group of animals require ∼22 sessions when a greater fraction of sessions are significantly above chance. However, we also observed considerable variability across animals in the number of recordings required to reach the desired level of performance (**Figure 2D**). Together these metrics indicated that mice are able to learn to detect vocalizations inside a textural background occurring at a random time.

**Figure 2.**
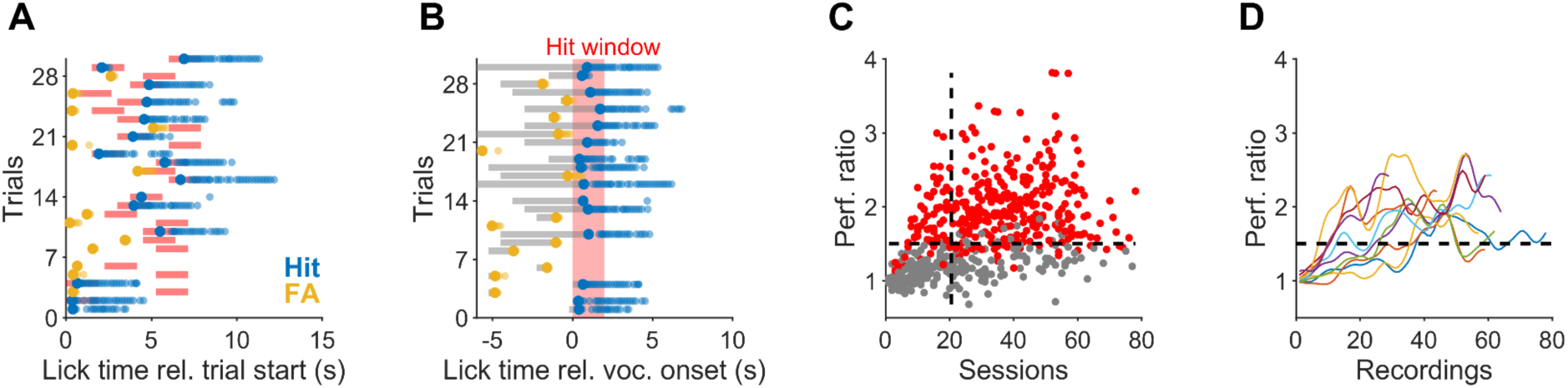
Mice learn to detect vocalizations embedded in textural noise. **A** After training the mice learned to respond to the onset of the vocalization. The mice continued to lick (blue dots) during and beyond the response window (red line). False alarms (orange dots) tended to occur close to sound onset with typically only one or a few licks. **B** Same data as in **A**, now aligned to vocalization onset (at 0, red line), showing that mice respond after the vocalization onset, indicating that they respond to the vocalization. **C** Progress in training over time as a function of performance ratio (real / shuffled hit rate) across all animals and recordings. Red dots indicate recordings where the animal performed above the criterion which was set to be 2×σ_rnd_ above the mean shuffled hit rate. Gray dots, on the other hand, correspond to recordings where the criterion was not met. Linear discriminant analysis revealed that animals tend to perform consistently above chance level after ∼22 sessions and once the performance ratio exceeds 1.5. **D** Same data as shown in (C), but session-binned for each animal individually. The black dashed line corresponds to the performance criterion derived in (C). Animals exhibit variability in terms of their learning rate but on average cross the performance criterion within ∼22 recordings (SD=12.43), confirming the analysis in **C**.

### Neural population response and pupil dynamics reflect the behavioral state

The task performance confirmed that the mice learned to actively listen for the vocalizations inside the textural background sound. Next, we checked whether the neural activity in the auditory cortex and the pupil diameter also reflected this engagement. Recordings were carried out both under active and passive listening conditions using identical acoustic stimuli. In the passive condition, the reward spout was moved away from the animal, hence, no lick response was registered and no reward was subsequently given.

Both the population response and pupil diameter exhibited an increase at the sound onset and elevated response following the initial adaptation period (**Figure 3A-B**). Neural and pupil data were normalized to their spontaneous average per neuron and animal, respectively (see Methods for details, and **Supplementary Figure 2** for alignment-based normalization). Active listening resulted in heightened responses for both metrics in comparison to passive listening (**Figure 3A**, aligned to sound onset: average normalized ΔFR_Active-Passive_=0.193; n=3621 matched units; Wilcoxon signed rank test, p=2.53×10^-95^, Cohen’s *d*=0.23; average normalized ΔPupil⌀_Active-Passive_=0.002; 2-way ANOVA, for behavioral state, df=1, F=303.9, p=4.74×10^-68^, Cohen’s *d*=0.01; **Figure 3B**, aligned to the vocalization onset: ΔFR_Active-Passive_=0.135, Wilcoxon signed rank test, p=7.36×10^-49^, Cohen’s *d*=0.18; ΔPupil⌀_Active-Passive_=0.003, F=140.79, p=1.80×10^-32^,Cohen’s *d*=0.02). When aligned to the time of the first lick (**Figure 3C**, active trials only), the population activity increases before the lick, while the pupil diameter remains unchanged. After the lick, the pupil diameter increases directly and substantially, indicating a close relation to the lick and the subsequent arousal (36).

**Figure 3.**
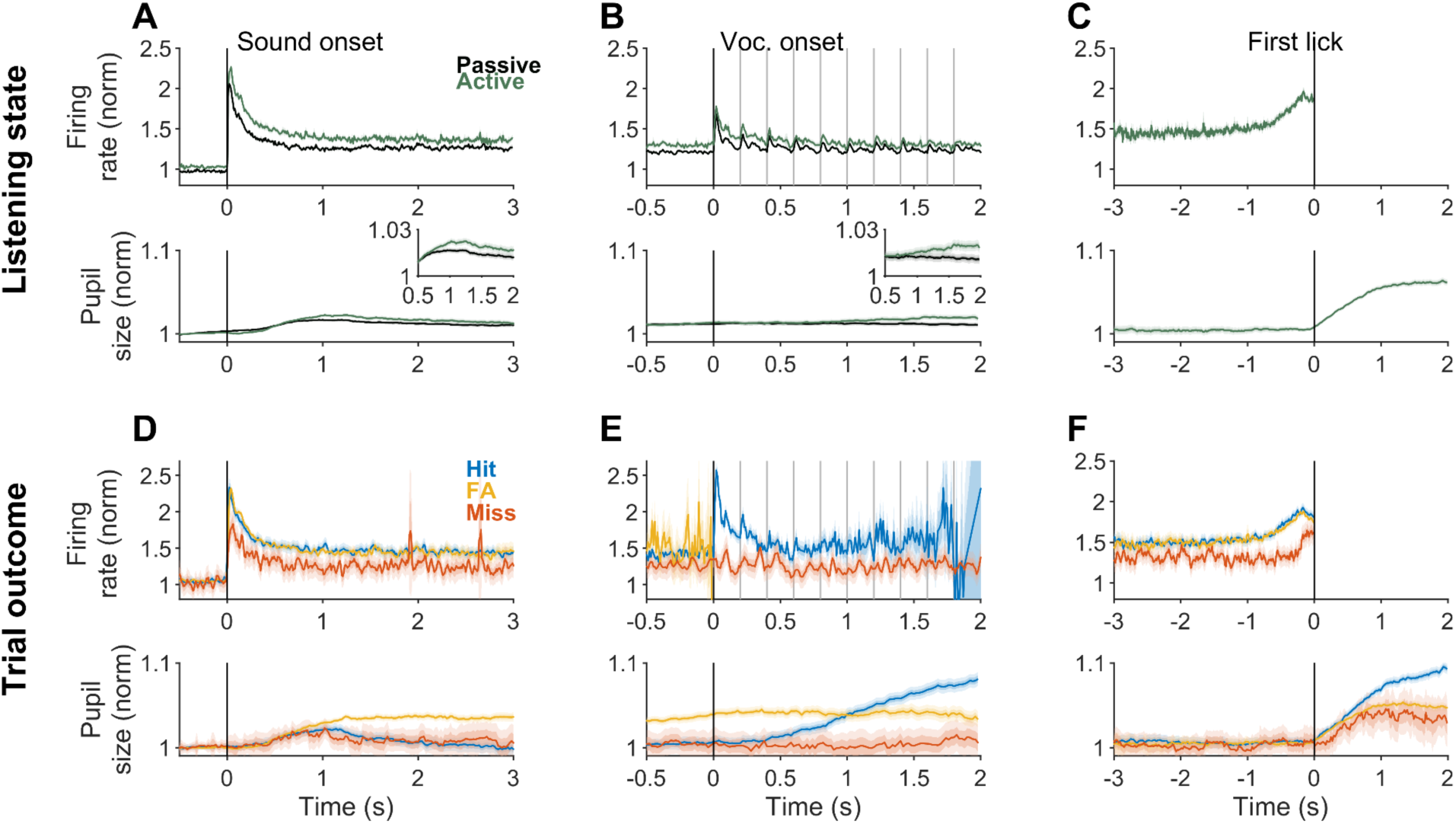
Neural population response and pupil diameter reflect behavioral variables reliably. Normalized firing rate (top panels) and pupil size (bottom panels) in active (n=12333 trials) and passive (n=19238 trials) listening conditions (A-C), and for different trial outcomes (D-F), aligned to sound onset (left column), vocalization onset (middle column), and first lick time (right column). **A** Active engagement in the task is characterized by elevated average firing rate as well as larger pupil diameters both when the responses are aligned to sound onset from trials where no vocalizations occurred before 3 s. The inset shows a zoomed vertical axis. **B** Same as (A), aligned to vocalization onset. **C** Neural population response shows a ramping prior to the first lick, which may be indicative of cognitive processes leading up to the decision to lick. **D** Sound-evoked population response was higher in hit and FA trials than in miss trials. Pupil diameter remained elevated in FA trials, likely reflecting the arousal after early licks in these trials. **E** Similar to (D), vocalization-evoked neural responses were stronger in hit trials compared to miss trials. Pupil dynamics primarily reflect different time points when the animal starts licking the spout. In FA trials licking starts before vocalization onset, in hit trials ∼0.5 s after, and in miss trials, the lick happens after vocalization offset. **F** While neural responses show ramping up prior to the first lick (top), pupil diameter increases after the lick reflecting arousal, which is heightened by reward in hit trials (blue). Neural and pupil data normalized to spontaneous averages per cell and animal, respectively (see Methods for Details and Supplementary Figure 2 for alignment-based normalization).

Next, we analyzed the data separately for different trial outcomes in active sessions. Similar to the differences observed between active and passive conditions, in trials where the animals were more engaged with the task (Hit and FA) the average sound evoked firing rates were higher in comparison with Miss trials, i.e. when the animal responds after vocalization offset (**Figure 3D**, top row, Kruskal-Wallis test, p=5.16×10^-5^). This difference was even more substantial in the case of the response to the vocalizations, where in miss trials the response was nearly absent (**E,** top row, Wilcoxon signed rank test between hit and miss trials, p=3.85×10^-20^, Cohen’s d=0.10).

Changes in pupil diameter also exhibited dependence on the trial outcome, but the time course of pupil diameter differed from that of the population PSTHs in important ways. While changes in pupil diameter in response to sound onset displayed a similar magnitude distinction between trial outcomes (**Figure 3D**, bottom, 2-way ANOVA, for outcome types, df=2, F=1.46×10^3^, p=0), vocalization onset-aligned pupil responses point to a closer relationship between pupil diameter and motor behavior or its relation with arousal (**Figure 3E**, bottom, F=717.8, p=1.46×10^-311^). After the initial peak in pupil diameter following sound onset, pupil diameter in FA trials remained elevated, while on hit and miss trials the pupil size went back to pre-trial levels. This likely reflects arousal after licking early in FA trials (**Figure 3E**, bottom, yellow). The average reaction time of mice in hit trials was 0.56 s (relative to first vocalization onset), which coincides well with the timing of the rise in pupil diameter in the same trials (**Figure 3E**, bottom, blue), consistent with earlier studies (37,38). On miss trials, the animals fail to respond on time to the vocalization, which is also consistent with the absence of arousal and thus flat pupil response observed in these trials (**Figure 3E**, bottom, red). The relation with arousal is further corroborated by aligning the responses to the first lick: While magnitude differences of population response under different trial outcomes are preserved (**Figure 3F**, top), the increase in pupil diameter only starts *after* the lick (thus effectively ruling out motor preparation as a source) and is larger in hit trials, where the reward is likely to increase arousal (**Figure 3F**, bottom, 2-way ANOVA, for outcome types, df=2, F=3.61×10^3^, p<ε).

Taken together, these findings show that average neural population response and pupil dynamics can distinguish between behavioral states in our paradigm. Consistent with previous research on detection tasks in the context of statistically defined stimuli, we observe general adaptation in response rate but also an increase in neural response prior to decision-making (39,40), while the pupil dilation appears to primarily reflect arousal. The increase in population activity before the lick may reflect the integration of evidence about the presence of the vocalizations (39,40), a change in attention due to the detection of a target (similar to (41)), motor preparation (42) or non-aligned sensory responses. We consider the latter two unlikely, given the long build-up (compared to much shorter reaction times in mice, (43)) and similarity of the build-up between trial outcomes (Fig. 3F).

### Detectability and neural encoding of vocalizations are improved by continued exposure to texture sounds

Next, we investigated whether the detection performance improved *within* a trial if the animals could listen to the background statistics for longer, as demonstrated previously in human subjects (39). For this purpose, we used the time-resolved d’ measure, introduced previously (39). The analysis relates the fraction of hits, correct rejections and false alarms that occur in a particular time window relative to stimulus onset to compute d’ (see **Figure 4A** for illustration, and Methods for Details). In agreement with the above-mentioned study, we observed a significant increase in the detectability of vocalizations as the prior exposure to the textural noise grew longer in duration (**Figure 4B**; Kruskal-Wallis test, p=0.005), suggesting an adaptation to or learning of the stimulus statistics as a basis for more accurate target detection.

**Figure 4.**
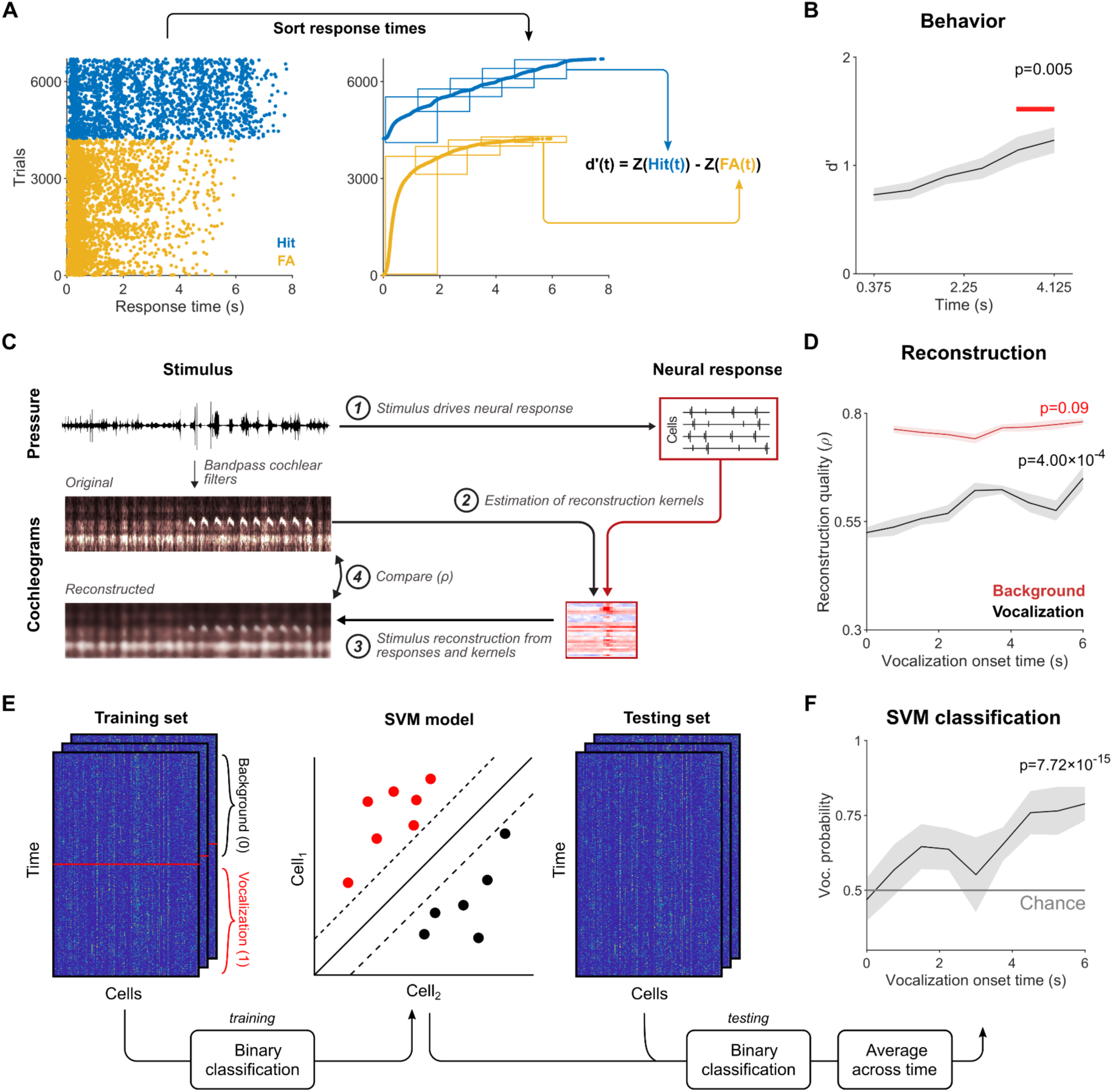
Detection performance and neural encoding of vocalizations improves with texture exposure. **A** We quantified detection performance using a time-resolved d’ measure, based on response times from all hit and FA trials (left) sorted in time (right). For each time bin, *t*, the proportion of hit and FA trials is computed. The difference between *Z(p)*, the inverse Gaussian cumulative distribution function for each type of trial, provides an estimate of d’ for each time bin. **B** The detection performance showed a significant increase over time into the trial (Kruskal-Wallis test, p=0.005, N=7 mice), i.e. with additional exposure to the texture. The red line indicates the time points after which d’ becomes significantly different from the first time bin. Gray hull indicates 1 SEM. **C** Reconstruction of sound stimuli was carried out by using a subset of stimulus-response pairs to estimate linear reconstruction filters. These filters were then used to reconstruct the original stimuli based on recorded neural responses. The quality of reconstructions was quantified by correlation between original and reconstructed spectrograms. **D** The reconstruction quality of vocalization (black) significantly increased relative to the vocalization onset time in the trial, while the reconstruction quality of the background (red) epoch did not significantly change (Kruskal-Wallis test, p-values shown in figure, hulls indicate 1 SEM). **E** A support vector machine (SVM) classifier was trained on a subset of data to perform the classification of the population activity vector as either vocalization (1) or background (0). The test set was then evaluated and binned across vocalization onset time bins as in **D**. **F** The SVM applied to the population vectors from the vocalization period showed a significant increase with vocalization onset time, consistent with both reconstruction analysis (D) and behavioral results (B). Flat line represents chance-level performance.

We hypothesized that the improvement in detectability might be related to an adaptation of the cortical responses to the background noise which would lead to a “pop-out” of the vocalization-related activity. Reconstruction of the presented sounds from the neural recordings allowed us to observe changes in the encoded information over time and has been used to demonstrate adaptation to background noise in human subjects (44). Using a similar approach, we reconstructed sound cochleograms by estimating linear filters from the activity of acoustically responsive cells (see **Figure 4C** for an illustration of this process, see Methods for details; (45,46)). Briefly, neural responses used for reconstructing the sounds were pooled across animals and selected (395 out of 1353 neurons) based on the signal-to-noise (SNR) of their spectrotemporal receptive fields (STRF). Data from passive listening sessions were used to avoid interference from licking artifacts. Reconstruction quality was defined as the correlation between the original and reconstructed cochleograms.

We divided the set of stimuli according to vocalization onset times and quantified the quality of reconstruction for vocalization epochs. In agreement with the behavioral data above, the vocalization epochs were reconstructed progressively better as the duration of exposure to the textural background sound increased (**Figure 4D**; Kruskal-Wallis test, p=4.00×10^-4^). While the background reconstruction appears in *absolute* terms better than the vocalization reconstruction, this might be a consequence of the reconstruction kernels being trained on texture-only data (see Discussion).

To validate this outcome using an alternative decoding method, we trained a binary support vector machine (SVM) classifier to distinguish the population activity vector at each time point as originating either from vocalization or from background-only intervals (**Figure 4E**). Subsequently, we assessed the classifier’s accuracy in correctly identifying population activity during the vocalization epoch (spanning a total of 2 seconds) as a function of vocalization onset time (**Figure 4F)**. A two-way ANOVA was performed to examine the effect of time within the 2 second-long vocalization epoch and vocalization onset time of the trial. The analysis revealed a significant main effect of time within the vocalization epoch (F=1.64, p=4.66×10^-8^), reflecting the adaptation to the repeated presentation of vocalizations. Importantly, we also observed a stronger main effect of vocalization onset time (F=97.07, p=7.72×10^-159^): Analogous to the reconstruction analysis previously outlined, this approach also demonstrated a gradual improvement in classification accuracy as the duration of prior exposure to sound textures within a trial increased.

Time-dependent improvements in both detection performance and the neural coding of the vocalizations themselves suggest that the auditory system of the mouse can integrate information over a timescale of multiple seconds to optimize the detection of an acoustic target stimulus in noise. These results align with the conclusions from human uECoG data that showed an improved encoding of the acoustic features of speech following an adaptation period after a switch in acoustic statistics (44).

### Behavior and neural activity improve at high cross-frequency correlations

Previous studies have shown that the auditory system is sensitive to long-term statistical regularities present in sound textures (18,30). In particular, cross-frequency correlations have been shown to significantly improve the texture identification performance in human subjects, suggesting that this statistical dimension carries important information about natural sound textures. Having established that animals can benefit from prolonged exposure to the textures in our paradigm, we varied the strength of marginal, pairwise correlations between frequency bands while keeping the marginal statistics of sound envelopes the same. The latter ensured that vocalizations could not be detected more easily based on sound properties other than CFCs. As the primary auditory cortex has been suggested to be involved in the processing of cross-frequency modulations (47,48), it constituted a natural first target to study the sensitivity of the auditory system to the correlational structure of the sound textures. By probing CFCs we also step beyond the consideration of marginal statistics only (39).

At a behavioral level, we observed that the animals were able to detect vocalizations that were embedded in high CFC textures more readily than those embedded in low CFC sounds (**Figure 5A**). This was true when the data were analyzed at the recording and the individual animal level (at recording level, Wilcoxon signed rank test, n=112, p=4.64×10^-11^, Cohen’s *d*=0.78) at the animal level, Wilcoxon signed rank test, n=7, p=0.0156, Cohen’s *d*=1.48). The neural population exhibited significantly less adaptation to high CFC textures, when aligned to trial start (**Figure 5B**, norm. ΔFR_High-Low_=5.9%; n=2065 matched units; Wilcoxon signed rank test, p=1.79×10^-19^), although the effect size (Cohen’s *d*=0.066) was much smaller than one might have expected due to the increased variability of the time marginal. The correlation between neurons was also higher in high CFCs compared to low CFCs (0.008 vs 0.011, respectively, p=0.0047, Wilcoxon signed ranks test). Pupil diameter also exhibited small but consistent divergence between the two conditions after the initial dilation related to sound onset (**Figure 5B**; ΔPupil⌀_High-Low_=0.002, 2-way ANOVA, for CFC level, df=1, F=233.8, p=8.63×10^-53^, Cohen’s *d*=0.002), likely reflecting the larger fraction of trials in which a reward was received (see above).

**Figure 5.**
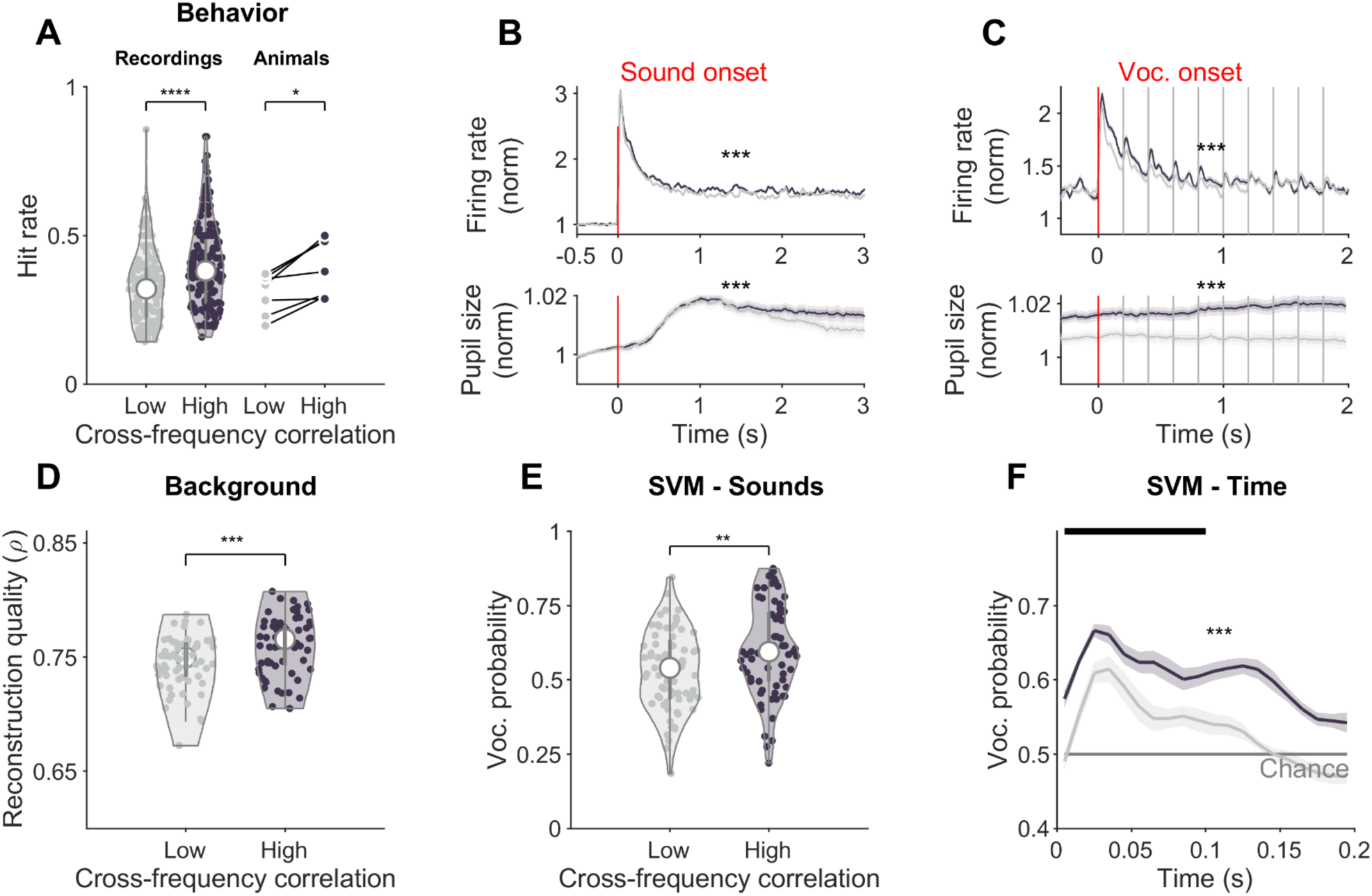
Detection of vocalizations is easier in the context of high cross-frequency correlations. **A** Behavioral performance across cross-frequency correlation (CFC) conditions was quantified as hit rates and analyzed at the recording and animal levels. At both levels, animals were significantly better at detecting vocalizations embedded in high CFC backgrounds than low CFC backgrounds. White dots on the violin plots correspond to the median. **B** The average PSTH (top) to the sound textures in active trials shows a small but significant difference, with high CFC adapting less than low CFC. The pupil response diverges after ∼1 s, indicating heightened arousal under high CFC conditions. Curves include only trials where vocalization has not yet been presented and FAs are only included up to the lick. **C** Average PSTH in response to vocalizations show a significantly greater response to high CFCs compared with low CFCs, consistent with comodulation masking studies. Again the pupil also dilates more in high CFC trials. Only responsive cells are included here. **D** Using stimulus reconstruction (see Figure 4C) indicates that high CFC textures can be reconstructed more clearly from the neural activity, suggesting that correlations allowed for a more faithful representation of the sound’s temporal structure. **E** Using SVM decoding (see Figure 4E) indicates that vocalizations were on average also represented more accurately in the context of high CFC textures (each dot is a stimulus presented). **F** Averaging the SVM decoding to detect population responses indicative of a vocalization (see Figure 4F) over stimuli and vocalizations (errorbar = 1 SEM) shows a clearly improved representation of vocalizations in the case of high CFCs (dark gray) in comparison with low CFC (light gray).

Population response aligned to vocalization onset exhibited a significantly larger response to vocalizations when they were embedded in high CFC textures (**Figure 5C** top; onset response (0-1 s): norm. ΔFR_High-Low_=10.1% (*d*=0.11, p=2.6×10^-30^); overall (0-2 s): ΔFR_High-Low_=5.7% (*d*=0.068, p=8.3×10^-14^); Wilcoxon signed rank test, n=1895 vocalization responsive cells, see Methods for details). The pupil response showed a significant and substantial increase in the high CFC as well (**Figure 5C**, bottom; average ΔPupil⌀_High-Low_=0.018, F=8.64×10^4^, p=0, Cohen’s *d*=0.18). If the neural response was normalized to the pre-vocalization period, the difference between low and high CFC was qualitatively and quantitatively similar as well as statistically significant (onset response (0-1 s): norm. ΔFR_High-Low_=5.4% (*d*=0.05, p=5.65×10^-15^); overall (0-2 s): ΔFR_High-Low_=5.9% (*d*=0.066, p=1.79×10^-19^)).

Together, this demonstrates that vocalization detection is aided by high CFCs of the background textures, as evidenced by improved behavior and higher vocalization related neural responses, in accordance with studies on comodulated noise (49). Importantly, an arousal increase, as indicated by the pupil dilation, is unlikely to lead to increased performance, but has been shown to reduce accuracy (50).

To relate this behavioral difference to the underlying neural responses, we again used reconstruction analysis as well as support vector machine (SVM) classification on neural data (as above in **Figure 4**). Reconstruction of background textures was more accurate for high CFC textures (**Figure 5D**; Wilcoxon signed rank test, n=64 stimuli per condition, excluding trials with no background-only epochs, p=1.85×10^-4^, Cohen’s *d*=0.63), likely related to the correlated stimulation of neurons with different frequency preference.

For decoding vocalizations we used SVM decoding (same as in Figure 4E/F), as reconstruction would conflate the background that was always present when vocalizations were presented. The SVM classified each time point as either vocalization (1) or no vocalization (0, i.e. background only), trained on all textures and then separately evaluated for high and low CFCs. Vocalizations could be decoded more accurately in the context of high CFC textures (**Figure 5E**; Wilcoxon signed rank test, n=72 textures per condition, p=0.011, Cohen’s *d*=0.498). Resolved across the time of a single vocalization (100 ms + 100 ms pause), the decoding of vocalizations was clearly more accurate in the case of high CFCs (**Figure 5F** means are averages across all 10 vocalization repetitions; 2-way ANOVA for factors CFC and time; for CFC, df=1, F=164.66, p=7.39×10^-28^; for time, df=199, F=2.81, p=5.04×10^-13^, Cohen’s *d*=0.93).

In summary, high CFCs lead to clearer neural responses that allow for improved representation of background stimuli and vocalization embedded in them. The latter could form the basis for the improved behavioral detection performance under high CFC’s, a phenomenon that has been described previously in human psychoacoustics under the term Comodulation Masking Release (49) which (see Discussion for further details).

### PV cells play a specific role in the processing of cross-frequency correlations

It has previously been suggested (reviewed in (51,52) that inhibitory interneurons could play an important role in highlighting a target sound in the context of an acoustic background, however, that this remained underexplored in the context of complex sounds. To study the contribution of inhibition to target detection in our paradigm involving complex textural sounds, we optogenetically manipulated the activity of inhibitory PV and SST interneurons. Optogenetic manipulation was performed in 30% of the trials. We titrated the light amplitude to limit the reduction in firing rate to ∼25% in cells that were not directly activated by light stimulation, in order to remain within a physiological range of network activity, i.e. to avoid completely silencing the response.

To analyze the outcomes of optogenetic manipulation, we classified recorded units as PV if their activity was significantly increased as a result of light stimulation (**Figure 6A**; (53)). While PV units with significant light response had spike shapes that were reminiscent of fast-spiking (FS) cells reported in the literature (sometimes also referred to as narrow spiking cells), we did not observe one-to-one correspondence between PV and FS cells **(Figure 6B)**. Therefore, we used these three partially overlapping groups of neurons to characterize the population response. The time course of sound-evoked response was similar across these cell types, but the average response of PV cells within the first 0.5s of sound presentation was significantly stronger than the other groups (Kruskal-Wallis test, p=0.001, **Figure 6C**). Light stimulation increased the sound-evoked response for PV cells (median=8.88%, Wilcoxon signed rank test, p=3.54×10^-5^), and decreased it for both FS cells (median =-3.84%, Wilcoxon signed rank test, p=0.0196) and non-FS cells (median=-11.85%, Wilcoxon signed rank test, p=1.16×10^-189^, **Figure 6D**).

**Figure 6.**
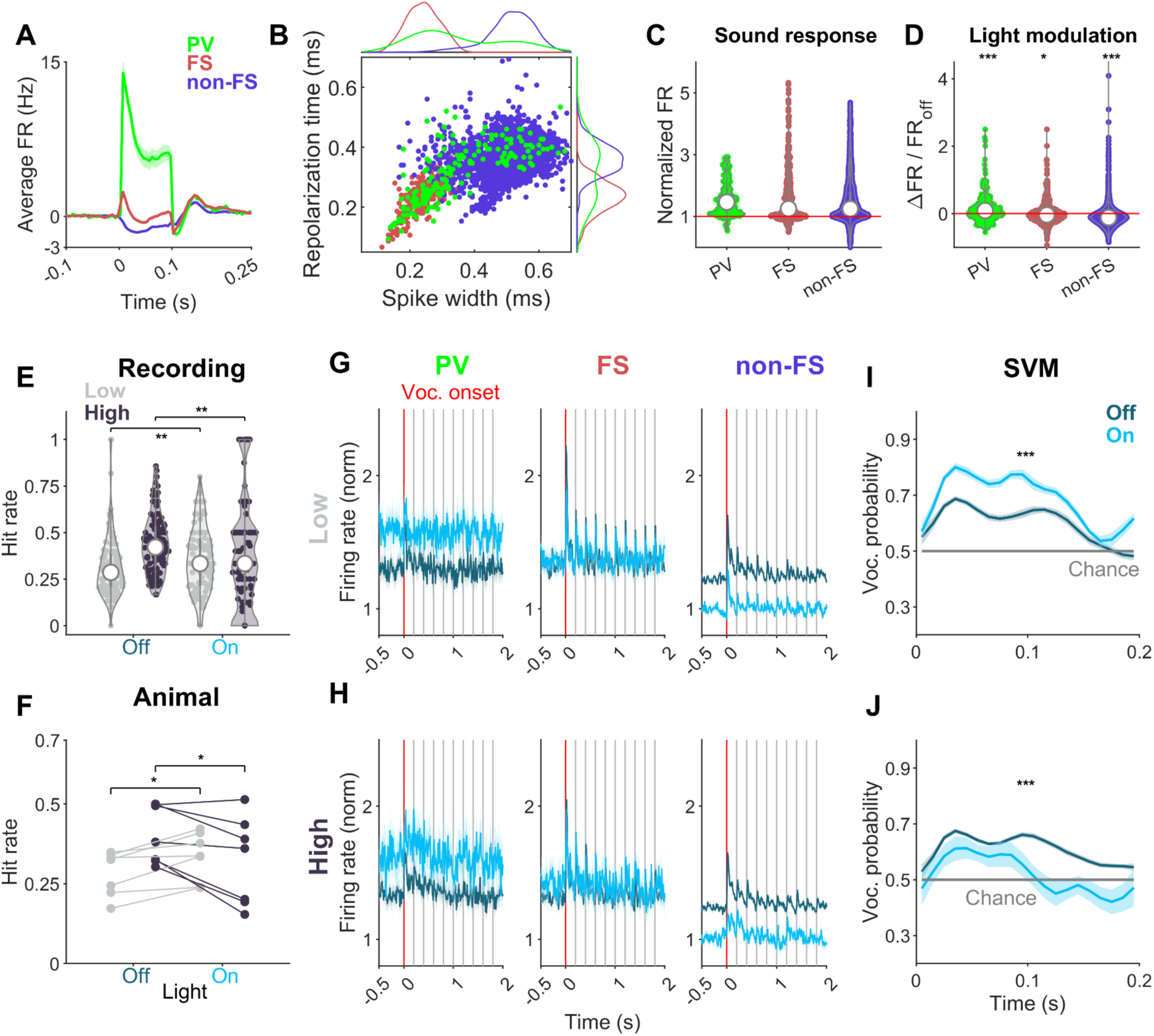
Optogenetic manipulation of PV cells has an asymmetric effect on behavior and neural encoding of vocalizations in different acoustic textures. **A** Average firing rate change in parvalbumin interneurons (PV, green; n=182), fast-spiking, (FS, red, n=439), and non-FS cells (violet, n=3741) in opto-tagging sessions. **B** Distribution of spike parameters shows PV cells tend to have narrower spike widths, though spike parameters do not uniquely identify PV cells. **C** All cell groups show significant sound-evoked responses relative to spontaneous firing rate (red), within 0.5s of texture onset. **D** Optical stimulation led to a significant increase in average sound-evoked response in PV cells, while FS and non-FS cells were inhibited to different extents. **E/F** Hit rates observed for low and high CFC under the light on and off conditions. Behavioral performance degrades under high CFC conditions but improves under low CFC conditions. Points correspond to different recordings (**E**, n=189) and animals (**F**, n=7). **G/H** Vocalization-onset-aligned PSTH (normalized to pre-sound activity) grouped according to cell types and background texture conditions (top/bottom row: low vs. high CFC; Light on trials: light blue, Light off trials: dark blue). Vertical gray lines indicate successive repetitions of vocalizations. Coarse features of PSTHs such as steady-state response level reflect the light reaction of each cell type shown in **D**. **I/J** We used the same SVM classifier as in Fig. 5E/F. The probability of being classified as a vocalization epoch was averaged over multiple presentations of vocalizations to yield a single time series for each condition. Despite having similar PSTH traces, the vocalization classification reflected the asymmetric effect of optical stimulation on behavioral performance (E/F). Gray line indicates change level performance. Error hulls represent 1 SEM.

The effect of optogenetic manipulation of PV was found to have opposing effects, depending on the type of background texture (**Figure 6E-F**). For high CFC sounds, activating PV cells impaired the behavioral performance (Figure 6E, at the recording level, Wilcoxon signed rank test, p=0.0025; **Figure 6F**, at the animal level, Wilcoxon signed rank test, p=0.0313), while conversely performance under low CFC sounds was significantly improved both at the animal level (**Figure 6F**, Wilcoxon signed rank test, p=0.0156) and the recording level (**Figure 6E**, Wilcoxon signed rank test, p=0.0062) as a result of this manipulation. Coarse features of the cell-type specific PSTHs were comparable across low and high CFC conditions, suggesting that the behavioral outcomes were not captured in any straightforward manner by population averages (**Figure 6G**). Therefore, to better understand the changes in the information content of neural responses, we used the SVM classifier to analyze the neural responses during the passive listening condition. As before, SVM evaluations mirrored the behavioral results analyzed at the animal level, where light stimulation improved the detection performance under low CFC background (2-way ANOVA; for light state, df=1, F=221.62, p=3.54×10^-34^; for time, df=199, F=3.99, p=2.63×10^-21^) and impaired the performance under high CFC condition (2-way ANOVA; for light state, df=1, F=89.55, p<𝛜; for time, df=199, F=1.62, p=3×10^-4^, **Figure 6H, I**).

Although the asymmetric effect of PV stimulation suggests a specific role played by PV interneurons in the processing of textures with high CFC, the outcome may still be due to the nonspecific effect of inhibition itself, rather than inhibition by PV cells per se. To evaluate this, we performed the same experiments on mice that expressed ChR2 in SST interneurons. While analyses involving no optogenetic stimulation trials showed comparable results (**Supplementary Figure 3C,F**), at neither behavioral nor animal levels, the SST dataset showed any asymmetric behavioral effect of optogenetic activation observed in the PV dataset (**Supplementary Figure 3D,E**). As was the case in the collection of the PV dataset, we titrated the intensity of the light to achieve a 25% reduction in population average response (compare to **Figure 6G** for non-FS cells) and used the maximum light intensity when the optical titration results were not conclusive. While this strategy led to effective control over population response in the PV dataset (Figure 6G, right column), we observed that the optical stimulation of SST cells (median=1.33%, Wilcoxon signed rank test, p=0.677) led to a weaker but significant inactivation of FS (median=-2.93%, Wilcoxon signed rank test, p=3.31×10^-5^) and non-FS cells (median=-6.93%, Wilcoxon signed rank test, p=1.54×10^-51^) (**Supplementary Figure 3B**). Therefore, even though differences between SST and PV datasets are strongly suggestive of cell-type specific roles for PV interneurons, we cannot fully exclude the possibility that the effect may be due to the overall level of inhibition.

### Neural coding transitions from samples to statistics within the trial

Lastly, we analyzed whether the neural responses contain more information about the specific realization or its long-term statistics, and in particular the contribution of different cell-types. As the statistical estimation appears to improve over time in a given trial, we hypothesized that the neural coding would transition from the specific realization to textural statistics over time. Extending an analysis in the visual system (54), we used a nested ANOVA to partition the total variance of neural responses into texture class and realization components: Texture classes were defined by their marginal and correlation statistics, while realizations corresponded to different segments of the synthesized sound clip belonging to a given texture class. Nested ANOVA can model the hierarchical relationship between realization and texture classes, and allows us to evaluate how sensitive the cells are to these features of the sound textures. Our goal in this analysis was two-fold: 1) how the explained variance (VE) for each component changed over time during the adaptation period, and 2) how the length of the analysis window affected the VE.

In order to address this question in a principled manner, we used moving average responses and performed nested ANOVA for each cell and time point, using different window sizes, which varied from 30 ms to 1000 ms in length **(Figure 7A**). We performed nested ANOVA separately on the three groups of cells (PV, FS, non-FS) introduced earlier. We then averaged the VE across cells to obtain the VE for the respective cell type population. While the population VE remained constant over the time window analyzed here (**Figure 7B-D**, left column), extending the sliding window length increased the texture class VE without improving the realization VE (**Figure 7B-D**, right column). With smaller sliding window averaging, VE by realization was greater than that by texture class for all cell types. However, when the responses were averaged over a longer period of time, the VE due to texture class increased significantly. Two-way ANOVA with cell type and window lengths used as factors for variance components identified cell type to be a significant factor for realization VE (two-way ANOVA, F=38.81, p=1.76×10^-17^), but not for texture class VE (two-way ANOVA, F=1.99, p=0.136). Window length had a significant effect for both sources of variance (two-way ANOVA, F=362.1, p<𝛜, for texture VE; F=66.35, p=4.07×10^-55^ for realization VE). These results show that the sensitivity of the population response in the auditory cortex varies based on the timescale of the analysis used and highlight the importance of the choices that are made in analyzing the sound responses where the temporal aspect is very critical. They also suggest that primary auditory cortical cells may be using temporal multiplexing when representing the noise, instead of converging on time-averaged statistics alone (55).

**Figure 7.**
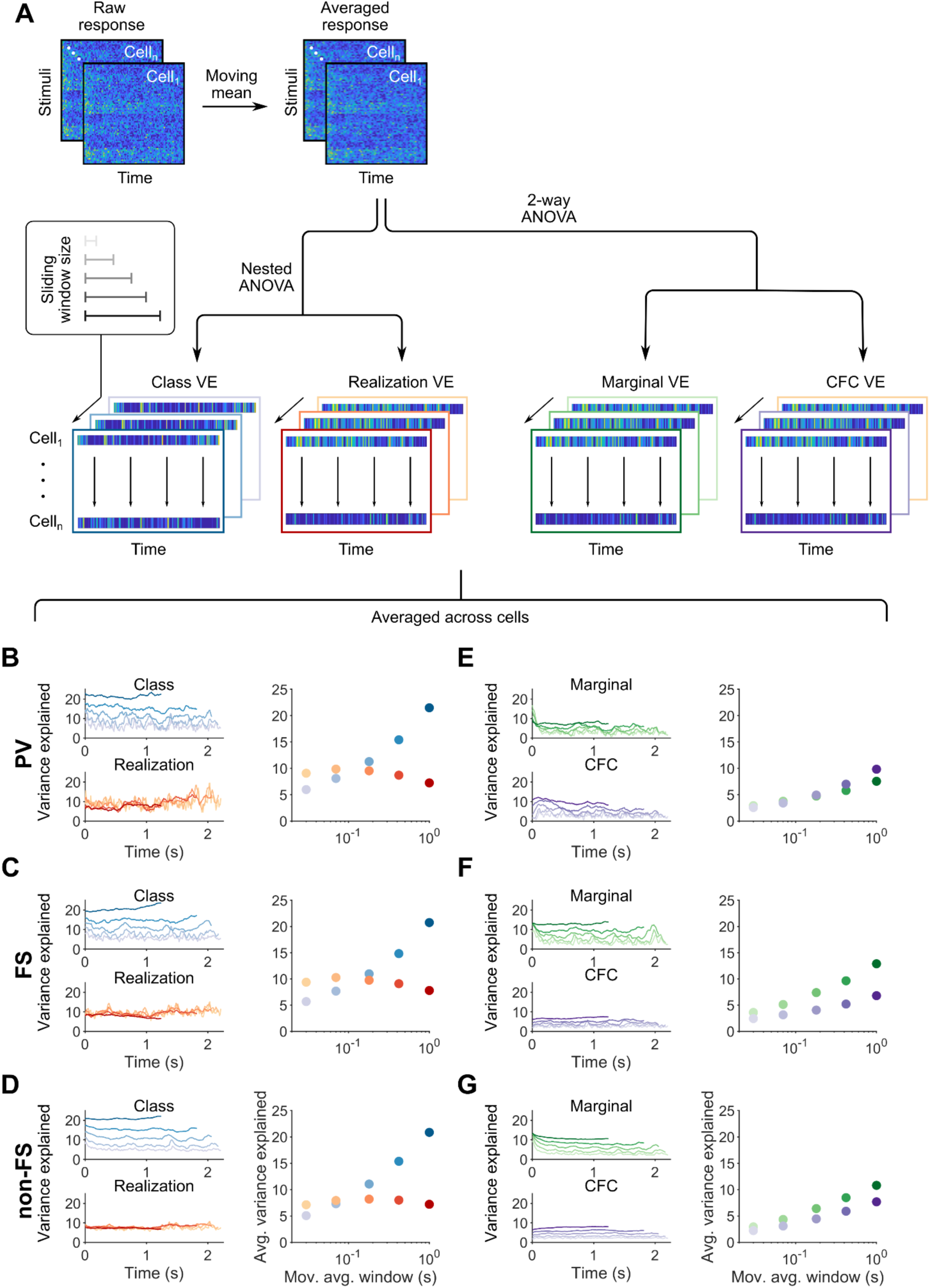
Sensitivity of different cell types to sound texture features. **A** The total variance of a cell’s response to sound textures was partitioned into contributions from different variables defining background sounds. We first averaged the cell’s response over time (first 2.25 s of texture only) using five logarithmically spaced, sliding window lengths. The averaged response of the cell was analyzed using either nested ANOVA (B-D) or two-way ANOVA at each time point. Variance explained (VE) was averaged across cells to evaluate change in VE over time as the animal listened to the sound, and to understand the effect of moving window size. **B-D** For shorter window lengths, realization VE exceeded class VE, whereas for longer windows, the opposite trend was observed. Nested ANOVA was conducted to partition the variability attributed to texture class and realization. The analysis was performed separately for PV (B; n=49), FS (C; n=192), and non-FS (D; n=1081) cells. All cell groups exhibited an increase in explained variance (VE) concerning texture class as window lengths increased, while their time courses remained relatively constant over the 2.25 s of response analyzed (left). To assess the influence of window length on variance component magnitudes, we calculated the average for each component over time (right). Notably, realization VE (yellow-red) displayed significant differences across cell types, whereas texture class VE (blues) did not (see text for statistics). Additionally, window length significantly affected both components. **E-G** Impact of marginal and CFC statistics on responses from various cell types. Consistent with B-D, we observed a progressive increase in VE by all cell types as the window size increased (2-way ANOVA). Both window size and cell type significantly affected the VE of both types of texture statistics, highlighting differences among the features that different cell types are sensitive to.

To further characterize the response variability, we performed two-way ANOVA using marginal and CFC statistics as factors, because these variables are not independent of one another (hence, not nested). Similar to the analysis above, the time course of VE by each of these factors did not show a strong pattern within the window analyzed here (**Figure 7E-G**, left column). However, in agreement with the previous analysis, average VE by both marginal and CFC components increased with longer moving average window lengths (**Figure 7E-G**, right column). Interestingly, both cell type (two-way ANOVA, for marginal statistics, F=11.73, p=8.19×10^-6^; for CFC statistics, F=4.74, p=0.0088) and window size (two-way ANOVA, for marginal statistics, F=114.26, p=2.36×10^-94^; for CFC statistics, F=63.32, p<𝛜) significantly affected the amount of VE by both types of texture statistics, suggesting some level of specialization in the type of statistics represented by different groups of cells.

## Discussion

We investigated the neural encoding of vocalizations in the context of textural background sounds in the auditory cortex of behaving mice. Both the behavioral performance and the corresponding neural encoding of the vocalization improved with exposure to the textural stimulus in individual trials, indicating a rapid, within trial adaptation to the textural statistics. Sound textures characterized by higher correlations across frequency channels (CFC) enabled the mice to more easily detect the vocalizations compared to the context of low CFC textures. Activation of parvalbumin (PV) interneurons exhibited an asymmetrical impact on behavioral performance for high and low CFCs, which was mimicked by the fidelity of the vocalizations’ neural activity, suggesting a specific role for PV cells in the processing of spectrotemporal correlations. These results contribute to our understanding of the neural basis for the well-documented ability of humans to comprehend conversations under adverse acoustic conditions, and in particular recognize and deal with sounds whose structure is only constrained statistically, such as many naturally occurring environmental sounds.

### Behavioral performance and adaptation to sound textures

Sound textures are characterized by their stable long-term statistics, and initial studies have shown that humans can utilize and estimate different types of statistics (18,30). These studies found that sampling a sound texture longer enhances the ability to distinguish between different texture classes, while at the same time compromising the ability to distinguish samples drawn from the same statistics, consistent with a neural encoding of statistics. Similarly, longer exposure to sound textures improved the ability to detect changes in statistics (39,56), phoneme recognition (44), and vowel discrimination (57). The process of estimation of statistical properties, which is believed to underlie these findings, occurs over multiple seconds, requires no conscious effort, and depends on certain stimulus features such as stationarity (58). In our study, texture statistics were changed on every trial, and thus also required the animals to adapt to these statistics within a given trial to optimize their performance. We find that the performance in detecting a vocalization inside texture noise also improves with exposure duration, and is thus consistent with an estimation process (Fig. 4). However, we did not observe plateauing in performance over time, previously observed in human studies (39). This discrepancy could be explained by task difficulty, where integration over longer periods might remain advantageous for more demanding tasks.

### Neural adaptation to and the encoding of sound textures

The neural encoding of sound textures has not received a lot of attention in previous studies, in particular not in the context of behavior and the detection of embedded target sounds, despite its paramount importance in real-world hearing. Here, we demonstrate that the encoding of vocalizations improves with time into the trial/stimulus, which we quantify through SVM decoding and stimulus reconstruction, respectively, while the quality of the encoding of the background texture remains unchanged (**Figure 4**). From previous human and neural modeling studies (11,44), we would have expected to find a reduced fidelity of encoding of the background texture, in conjunction with an improved encoding of the foreground. While we find the generally expected adaptation in firing rate (5) to the texture stimulus in line with (44), we do not find a specific buildup of invariance against the background. Within the limits of the current task, this is hence not directly in line with the general idea that highlighting of relevant information - which we find to be the case - is necessarily accompanied with a reduction of information about background sounds (46), at least not in mouse A1. Higher level areas might show different degrees and properties of adaptation to different stimulus contexts (59). In addition to the species difference to (44), in our case the reconstruction model was not trained on clean speech neural data (not available in our case), but on textural noise responses, which probably improved the overall reconstruction quality for textures at the expense of vocalizations.

On the other hand, the observation that vocalization reconstructions improved over time is consistent with the jointly observed, improved behavioral detection performance, as well as with previous studies into the effect of prolonged exposure to masking noise that showed a stronger signal-evoked response (48), improved reconstruction of the speech signal (44) and improved detection and recognition of foreground targets in the context of textural noise (22).

The most fundamental form of adaptation that occurs in the sensory systems is firing rate adaptation, where the firing rate of neurons to an ongoing stimulus is reduced over time (60–62). Across the auditory system the time-scale of this adaptation ranges from 10s of milliseconds (63,64) to 100s of milliseconds (Fig. 3/5), but does not extend to multiple seconds as observed presently in the within trial performance improvement. Further, response magnitude and subsequent adaptation to the sound in the primary auditory cortex are known to be sensitive to variables such as task engagement (65–67) and attention (68). In agreement with previous studies, we observed differences in the magnitude of sound-evoked responses between active and passive listening conditions, as well as trial outcomes. These differences were also observable in pupil diameter over time, and likely reflect the aggregate nature of metrics like average population response and pupil dilation (69).

Further, we observed that A1 responses carry information about different features of the sound at different timescales (**Figure 7**), consistent with the dual timescale encoding of sound observed in previous studies on the perception of sound textures (39,70). At faster timescales, the variance of all cell types analyzed here is better explained by the realization (i.e. fine temporal details of the sound clip), whereas at the scale of multiple hundred milliseconds, neural responses were better explained by the texture identity, which is a higher-level, time-averaged feature of the sound. This is consistent with the idea that the neural responses may carry information at different timescales, a phenomenon known as temporal multiplexing (55,70–73). The existence of neural codes on different timescales in A1 can in turn underlie the duration dependence of texture class and exemplar differentiation (30), if suitably integrated by higher order areas.

Previous work on the sensitivity of different auditory regions to texture statistics in the anesthetized rat has shown that while the inferior colliculus (IC) is remarkably sensitive to most classes of statistics in the McDermott and Simoncelli model (18), A1 does not reliably respond to changes introduced to ongoing texture statistics (24,26). In contrast, we find in the awake mouse that (in particular for long time windows) texture type can explain ∼20% of the variance (**Figure 7**), and substantially more than different realizations with similar statistics. On the one hand his could be due to the difference in state, i.e. awake vs. anesthetized, or species, i.e mouse vs. rat. In addition, the stimulus design also differed, where Peng et al. studied changes between different textures, while we focussed on the neural encoding of textures in separate trials. Overall, the strong adaptation of cells in A1 to ongoing background sounds (20,23,44) could also limit the ability to decode texture type in both studies, which, however, might be a reflection of a noise reduction strategy in the cortex. In this context, the constant ability to reconstruct the spectrogram over time (Fig. 4D) is puzzling, and may suggest a greater coding efficiency with longer presentation duration.

A recent, highly insightful study on active sound detection in noise (29), closely linking experimental data to a normative model. Their primary manipulation - switches between low and high contrast textures - led to two different time-courses of adaptation in both behavioral performance and neuronal sensitivity. However, their contrast change does not map directly onto our low/high CFC paradigm, as only low CFC textures are used, limiting a direct comparison of the result. In particular, differences in integration windows (max. 1 vs. 6 seconds), timing (expected vs. unexpected context duration) and behavioral performance measure (percent correct vs. d’) complicate a direct comparison. Framing the present paradigm in their normative framework extended to include correlations and longer timescales would be required to fully compare and reconcile the observed adaptation dynamics between the two paradigms.

### Sensitivity to cross-frequency correlations

On the behavioral level, we observed higher detection performance in high CFC textures (**Figure 5**), which aligned with results on the neuronal level, where we observed slightly stronger adaptation to low CFC stimuli and significantly stronger vocalization-evoked neural response in a high CFC context. The presently observed effect size in the neural response is small, but comparable to previous studies (47,74), which also highlighted a dependence on the relative level of masker and target, which was set to a fixed relation in the present study. The improved performance for high CFC stimuli cannot be explained by faster convergence statistical properties for these stimuli, which we checked directly on the level of the stimuli for different excerpt durations (see **Supplementary Figure 4**). More importantly, we find a better encoding of vocalizations for high CFC textures (**Figure 5E/F**), and a strong, asymmetric effect of increased PV activity on the encoding of vocalizations for high vs. low CFC backgrounds (**Figure 6**). This generally aligns with previous work that demonstrated that the average frequency bandwidth of neurons across successive stages of auditory processing tends to increase from auditory nerve fibers to A1 (75). Due to this feature, cells in the cortex might be well suited for integrating information across different frequency bands.

Given the pronounced comodulation in the presently used high CFC sounds, the behavioral and encoding improvements could relate to comodulation masking release (CMR), which describes the facilitation of narrow-band sound detection when different frequency bands in the background noise are comodulated in level (32). The latter is considered to provide a spectral cue for grouping different parts of the spectrogram which the auditory system could use to segregate non-comodulated target sounds from the background (76–78). Previous studies have shown that there is evidence on the level of neural responses on multiple levels of the auditory system, ranging from the cochlear nucleus (79) to the auditory cortex (47,48,80,81). While a rich literature exists on CMR in humans, the limited number of studies in animals have typically not studied behavior and neural encoding together, in particular not in conjunction with cell-type specific modulation as done here (25,47,48,74,81,82), which has left the concrete neural basis of CMR unresolved. Importantly though, the present study only has limited relevance to CMR: while the behavioral and neural improvement in the high CFC condition is closely related to CMR, our main results, i.e. (i) integration of statistics over time, and (ii) role of PV interneurons, are either general to both low and high CFC (i) or contribute constructively to the low CFC (i.e. non CMR) condition (ii). Therefore, they contribute to the understanding of listening in noise, even in conditions when no comodulation of close or distance flanking bands exists.

Another factor contributing to the improved performance in high CFC background could be related to “dip listening”, i.e. the ability to catch acoustic “glimpses”, when the level of noise momentarily drops (83). In the high CFC case, the SNR of the vocalization relative to the background fluctuated more, and high SNR moments could have been used to improve detection performance. It is, however, noteworthy that modulated noise does not always aid the recognition of signals in a complex auditory scene, as it can also transiently capture the attention and thus interfere with target recognition (84,85).

### The role of inhibitory neurons in processing background noise

We characterized the role that PV, SST, FS and non-FS cells may play in the processing of naturalistic noise in the primary auditory cortex of the mouse. Identifying cell-types is crucial for understanding the underlying processing, e.g. hearing in complex auditory scenes with competing sound sources (15,48,86). As there remains debate on whether opto-tagging (parvalbumin, PV, and somatostatin, SST) or spike parameters (87–89) is the best approach for classification of cell-types, we opted for using both in parallel (**Figure 6**).

While the cellular basis of hearing in complex auditory scenes is only starting to be explored, various fundamental aspects of it such as surround suppression, adaptation to repeated sounds, and the role played in attentional control have been studied previously (see (52) for a detailed review). These particular aspects of sound processing are important in real-life audition because natural sound sources usually differ from basic sounds used in auditory experiments by wider bandwidths, more spectral overlap and more diverse temporal dynamics. This poses a challenging problem which could require filtering out interfering noise and selectively attending to relevant sounds. It has been shown (14) that while wideband noise drives both SST and PV interneurons, only inactivating SST cells leads to a change in surround suppression observed in excitatory cells. Stimulus history modulates the inhibitory effect of SST cells, while PV cells were shown to provide the same amount of inhibition regardless of the stimulus history (9).

At first glance it may seem that this contrasts with our results, namely that only the activation of PV cells influenced the behavioral and the neural activity. However, upon close inspection the results are compatible: the aforementioned results(14) emphasize the role of SST cells in integrating *broad-band noise*, and conversely a lack of a role of PV cells for the same task. This aligns with our result, as we find a beneficial role for integrating *narrow-band* noise, as in the low CFC case (see **Figure 8** for a graphical summary). Previous studies have directly demonstrated differences in tuning width between PV and SST cells, with an awake study showing a wider integration in SST cells (13). It should be noted that a study under urethane anesthesia found the opposite result (90), which has been argued to be potentially due to the strong attenuation of anesthesia of SST cell activity (91).

**Figure 8:**
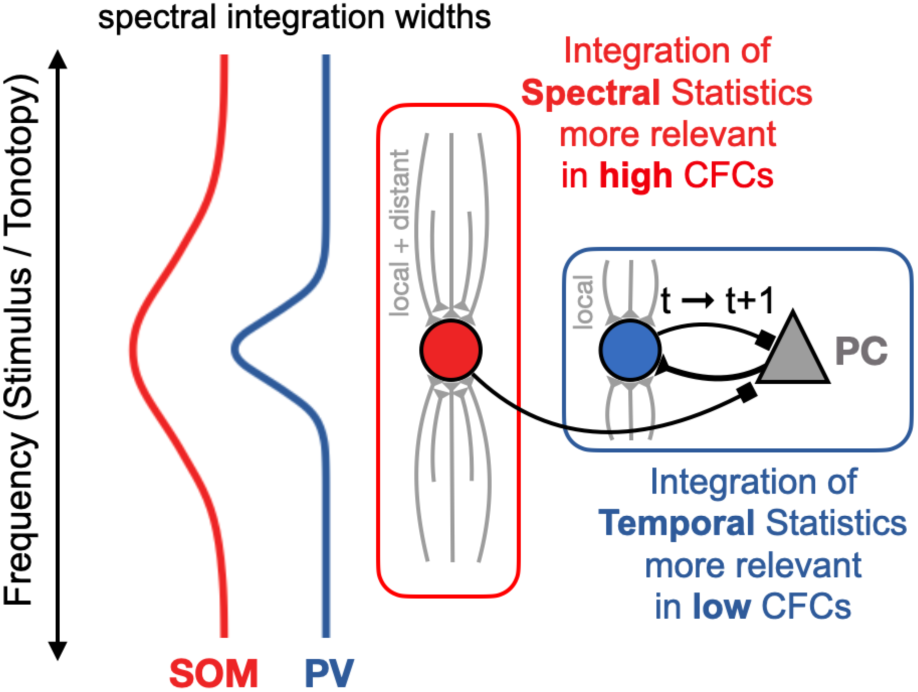
Putative roles of PV and SOM cells in the integration of spectral and temporal statistics. The activity of PV and SOM interneurons both modulate the activity of pyramidal cells in the cortex. However, PV and SOM interneurons differ in their spectral integration width, with SOM cells integrating more widely than PV cells (Kato et al. 2017). This would allow SOM cells to contribute to the integration of spectral statistics, e.g. in the case of high CFCs (Lakunina et al. 2020). As presently shown, the activation of PV cells leads to improved encoding of vocalization in low CFCs (and vice versa). Together with the known reciprocal connections between PV and PC, this suggests that they contribute more to narrowband, temporal integration of statistics. Gray connections indicate a mixture of intracortical and thalamocortical inputs.

As noted above, high CFC sounds had stronger but randomly timed amplitude modulation, in contrast to the relatively flat time course of low CFC sounds. It makes sense that this might not have been detectable in the aforementioned study, where narrow-band context was not assessed. Conversely, in the *broad-band noise* case, we find PVs to have a detrimental effect, which we interpret as a shift in the balance of activation between different cell-types, which might interfere with the SST’s beneficial effect on broad-band integration. To our surprise, direct modulation of SST neurons did not lead to a behavioral difference in either of the CFC conditions, which raises the question whether the results from (14) generalize to textural stimuli. The aforementioned study used basic white noise, while here we used naturalistic textural sounds with greater internal predictability, both in time and across frequencies. Consequently, the improved performance during PV stimulation in low CFC sounds suggests a more direct role in temporal coding, as in the low CFC case within-channel temporal predictability is greater, which aligns with previous findings on the role of PV cells in temporal coding (86). The local co-tuning of PV cells with excitatory cells has been suggested to reduce the noise in the neural activity (86), which presently translates to subtracting the recent, narrow-band noise. This can form the basis for improved signal-to-noise for the embedded vocalization, and thus provide the basis for the improvement in behavior and neural encoding. Slightly reduced adaptation and improved reconstruction quality for high CFC sounds suggest that in the context of our task, the primary role of PV mediated inhibition may be to shape the population response in order to serve something akin to temporal envelope tracking or neural entrainment (92–94).

### Limitations and future directions

We presently chose to increase the activity of PV/SST cells (similar to other studies; (95,96), in order to understand their influence on the encoding of textural sounds and subsequent behavioral decisions. Instead one could have silenced these cell-types, which would be more consistent with the classical lesioning approach that removes a part of a system to understand its functional contribution (97). While optogenetic modulation using light-activated Cl^-^ (GtACR, NpHR, etc.) or H^+^ (Arch) channels (see (98) for overview) would have allowed for this in a transient and reversible manner, it is not clear whether this would have been superior, as it risks moving the system outside of its physiological range (99) which complicates interpretation in highly interconnected systems (100). In our perturbational approach, we instead aimed to modulate the average activity of the non-PV/SST cells in a limited manner and thus keep the system close to its physiological range. Further, training of the SVM and reconstruction algorithms was always balanced for correlation, light stimulation, and vocalization base frequencies, and should therefore not have biased the quality of decoding towards either high or low CFCs. The observed, interpretable changes in behavior and neural activity for the high/low CFC stimuli seem to validate this approach, however, we would like to point out that it is inherently limited, as the modulation is not specific to individual cells, and depends on the penetration depth of the light. Future studies should improve on this technique by the use of more specific stimulation techniques such as Spatial Light Modulators (SLM) alongside optical imaging (as in e.g. (101,102)).

Optogenetic manipulation of PV and SST cells make a case for a specific role of PV-mediated inhibition in the processing of high CFC noise (see **Figure 6**). However, the behavioral difference between PV and SST may arise from different levels of inhibition achieved in each case. While we used identical light delivery in both cases, there are factors that could have limited the amount of inhibition in the SST case: (1) Physiologically, PV cells form inhibitory networks, wile SST cells avoid each other and form disinhibitory circuits with other interneuron types, which may limit the degree of inhibition achievable (103). (2) More PV cells tend to be located in superficial layers than SST (104), which simplifies their stimulation in particular in chronic experiments, where scarring may reduce the amount of light that penetrates deeply enough. Future studies utilizing more local light delivery or deeper penetration using 2-photon stimulation could further resolve this question.

The current recordings were targeted to the primary auditory cortex, and there is clearly a need to study the corresponding activity on different levels of the system, starting from the inferior colliculus to higher stages of processing, in particular the parietal cortex which has previously been demonstrated to be involved in the integration of sensory evidence (105). Chronic widefield and 2-photon optical imaging using more recent calcium indicators (106), which provide high temporal resolution and excellent signal-to-noise ratio, combined with targeted optical stimulation are likely the best techniques for further advancing this research, as they allow to resolve the spatial layout of the system with little to no undesired perturbation.

Lastly, while the present set of stimuli was already large, reaching the time limits of working with behaving mice, future studies should address additional dimensions in the space of stimulus statistics. First of all, we only studied two cross-frequency correlation levels, and resolving this more finely, or introducing other naturalistic CFC patterns would be very important. In addition, introducing different temporal correlation widths would be another interesting direction to demonstrate the involvement of PV interneurons in temporal integration. Each of these questions would have to be addressed in separate studies to obtain sufficient data for the specific stimulus contrast, comparable to the low/high contrast studied by (29).

## Methods

Data presented in this study are collected from 7 PV::ChR2 (4 males, 3 females; age at experiment mean = 10.2 weeks, range = 9-13 weeks) and 3 SST::ChR2 (3 males; age at experiment mean=10 weeks, range=9-12 weeks) mice obtained from F1 generation of crossing between either PV-Cre (JAX stock #017320) or SST-IRES-Cre (JAX stock #013044) with Ai32 (JAX stock #024109). 4 animals (2×PV::Ai32 and 2×SST::Ai32) were used for hearing ability tests (see below). Animals were housed in a reversed light/dark cycle environment (12h light/dark), inside the cages where food pellets were available *ad libitum*, while water access was limited to motivate animals to learn the behavioral task (see Details below in Animal Training). Most animals were housed in pairs separated by a perforated transparent plexiglass wall that allowed limited interaction between animals while protecting the implant from damage and allowing per-animal monitoring of water intake. All animal experiments were performed under animal experimentation project 2017-0041 approved by the Centrale Commissie Dierproeven (CCD) and supervised by the Animal Welfare Body of the Radboud University Medical Center.

### Surgery

Animals underwent two surgeries with a training period (6-8 weeks) between them. In the first surgery, a cement base holding a headpost was implanted for head-fixed training, while in the second surgery, the electrodes were implanted (see details below). For both surgeries, analgesic treatment began 1 day before the surgery and continued for 3 days post-surgery using oral intake of an analgesic diluted in water (Rimadyl, 5 mg per 100 ml). Animals had *ad libitum* access, with the concentration estimated based on their normal water intake. At the beginning of each surgery, mice received an additional dose of Rimadyl (5 mg/kg, s.c.) and in the second surgery, a dose of dexamethasone (2 mg/kg, s.c.) was used to reduce brain swelling.

In the first surgery, a plastic headpost for head fixation was implanted on the contralateral cranial surface of mice that were 6-8 weeks of age. Implantation was performed under isoflurane anesthesia (1.5-2%) while continuously monitoring breathing rate and body temperature. The animal was placed on a thermal blanket and the eyes were covered with eye ointment (Ophtosan, ASTFarma). After shaving the head and prior to making the incision, 0.1ml of Lidocaine/Bupivacaine was administered subcutaneously using a fine syringe needle. After removing the skin over the cranium, the bone was cleaned from tissue using hydrogen peroxide (H_2_O_2_, 3%). Next, a 3D-printed, plastic headpost (material: polycarbonate, Prusa) was attached to the cranial surface using dental cement (Super-Bond C&B Kit, Sun Medical), positioned approximately midway between bregma and lambda and close to the lateral ridge, leaning laterally at a ∼50° angle from the midline. In a subset of animals, a round glass plate was placed over the auditory cortex for intrinsic optical imaging of the auditory cortex (see below).

In the second surgery, the mice were implanted with electrode arrays in the primary auditory cortex (A1), whose location was estimated by either stereotactic coordinates (2.8 mm from Bregma, 5.9 mm laterally from sagittal suture; based on (107) alone or together with the help of intrinsic optical imaging (see below). A small craniotomy of ∼1×2 mm in size was opened and a small amount of biocompatible silicone gel (Dura-Get, Cambridge NeuroTech) was placed on the craniotomy to minimize brain swelling. Electrode arrays (see Details in the section Electrophysiology below) were inserted orthogonal to the brain surface with the prongs aligned along an axis that maximized the coverage of topographically organized frequency space (based on the orientation of area A1 in the map of (107)). A ground screw was attached contralaterally, anterior to the headpost, and connected to the ground cable of the recording probe using conductive epoxy (CW2400, Chemtronics).

### Intrinsic optical imaging

In a subset of animals (7/10), intrinsic optical imaging of the auditory cortex was performed to locate the primary auditory cortex and improve targeting area A1. During the implant surgery, the mouse was lightly anesthetized with isoflurane (∼1%). Pure tones and noise bursts were presented (70 dB SPL) as probe stimuli to activate the auditory cortex. The surface of the brain was monitored optically through a glass window (ø=3 mm, CS-3R, Harvard Apparatus) placed onto a thin layer of cement, centered on the region of the skull that was estimated to be above the auditory cortex based on stereotactic coordinates. The brain surface was monitored through the cement and skull using an sCMOS camera (CS2100M-USB, Thorlabs), receiving light from a combination of a 4x objective (N4X-PF, Nikon) and a tube lens (TTL-100A, Thorlabs). The recording area underneath the glass was first illuminated with a collimated LED (525 nm, M525L4, Thorlabs) to obtain a reference image emphasizing the blood vessels. During functional imaging, illumination was switched to 625 nm (M625L4, Thorlabs) and imaged at a rate of 20 fps. The focal plane was placed roughly 100 μm below the vessel level.

The acquired movies were analyzed by a sequence of normalizing and filtering steps: (1) reduce image resolution by averaging over 4x4 blocks, (2) normalize frames for differences in global illumination based on the average intensity outside the craniotomy, (3) baseline correct frames for each pixel by dividing them by the average intensity in the pre-stimulus period, (4) reduce the temporal sampling rate to 10 fps by averaging consecutive frames, (5) average over collected trials, and (6) perform spatial filtering by convolving with a 2D Gaussian kernel (SD = 2 pixels). The resulting response maps were then obtained at the minimal response after the stimulus onset, indicating the time of the largest activity-dependent change in blood flow, volume, and oxygenation in the observed brain regions (108). The resulting spatial pattern was then compared with previously established maps of the auditory cortex (107) to identify the most likely location of A1 in relation to the local vasculature.

### Electrophysiology

All recordings were performed using silicon probes featuring 64 channels (NeuroNexus) of various geometries (N_prongs_xN_sites_), specifically two 8x8 (A8x8-Edge-5mm-50-150-177, A8x8-Edge-5mm-100-200-177) and two 4x16 (A4x16-Poly2-5mm-23s-200-177, A4x16-2.5mm-tet/lin-300/125-333-121/177-PC). Data was digitized using a combination of 64 channel headstages (Intan Technologies) connected to an OpenEphys recording system (Open Ephys Inc.). The data at a sampling rate of 30 kS/s, band-pass filtered between 0.3-7.0 kHz, and then spike-sorted to obtain single and multiunit responses using autoSortC function included inside the code repository provided along with this article. Electrophysiological recordings were collected over a period of 4-8 weeks, depending on the signal quality. Electrodes were advanced weekly (in steps of 125-250 µm) to cover the entire depth of the cortex over the whole recording period. At the end of the experimental period, animals were disimplanted, and intact probes were cleaned (Tergazyme) and reused.

### Optogenetic stimulation

Cell-type specific manipulation of PV interneurons was carried out by photostimulating blue-light activated opsins (ChR2) 470 nm light was delivered using an LED light source (Optogenetics-LED, Prizmatix, maximum light power ∼10 mW). Dual fiber optic cannulas (manufactured by Prizmatix) were inserted into the custom-made implant case through tunnels on both sides that lead to the brain surface. The end of the optical fiber was located ∼1-1.5 mm above the brain surface. For determining the level of light on a given day, short recording sessions were carried out, where we delivered 100 ms light pulses of varying light intensities, separated by a pause of random duration (1-1.5 s). These sessions were used to opto-tag the cells that were responding directly to the photostimulation. Additionally, the power of the light delivered at each recording was titrated based on these sessions to induce a 25% decrease in population (with PV cells excluded) firing rate as a result of activation of PV interneurons. The goal of this titration procedure was to modulate the influence of PV interneurons in a controlled manner, but not completely silence the auditory cortex, which we expected would abolish the behavior (29). During the main experiment, light was provided at the same intensity as above, during a randomly selected but fixed set of trials. If the titration experiment failed to record the expected inverse relationship between the light intensity and population firing rate (often due to scar tissue that developed over time), the light intensity was set to the maximum possible by the device. The light was turned on at the same time as the sound with a ramp of intensity at the beginning and the end of the stimulation to reduce the stimulation artifact.

### Pupillometry

The left eye of head-fixed animals was recorded using a high-speed camera (GV-5240CP-NIR-GL, IDS Imaging Development Systems) under collimated infrared (740 nm) light illumination (WLS-22-A WheeLED, Mightex) delivered through a light guide. Images were captured at 50 fps and compressed to MPEG-4 files (quality ratio=0.5) before being further processed. Pupil diameter was estimated using a custom script utilizing pose-estimation tool, DeepLabCut (109). The model was trained to estimate positions of 6 points around the pupil perimeter as well as the center of the pupil. These points were then used to compute multiple radius and diameter estimates for each frame. To account for mislocalized points, the final pupil diameter was computed as the median of all measurements. Time points, when an animal blinked, were omitted from the analysis, as the marker values were not available for the corresponding frames.

### Facial motion analysis

We conducted a hearing ability test on untrained mice (n=4; 2×PV::ChR2 and 2×SST::ChR2) using the same setup as previously described. The mice were exposed to a randomized sequence of pure tones, while their facial expressions were recorded. The experiment involved presenting pure tones of five different frequencies (2, 4, 8, 16, and 32 kHz) and a silent stimulus as a control condition. Each tone was played for 2 seconds with a random pause between tones lasting from 4 to 6 seconds. The mice were exposed to each tone 50 times during the experiment. The videos were recorded and subsequently subjected to singular value decomposition to reduce their dimensionality, as detailed elsewhere (38). To estimate the hearing ability of the mice for each frequency, we identified the peak of the average motion energy time series from the first motion component (**Supplementary Figure 5**). We performed only one session per animal to prevent habituation to the experimental context.

### Acoustic stimuli

Acoustic stimuli consisted of a background texture and repeated, synthetic mouse vocalizations. Sound textures were synthesized using the Sound Texture Synthesis Toolbox (18) adapted to our particular use case. To generate sounds that vary mainly along a limited set of statistical dimensions of interest, we first measured the respective statistics in a set of naturally occurring sounds provided by (18). Using measurements from different natural sounds, we synthesized artificial sound textures that had specific subband means ("frequency marginals") and variances as well as particular cross-frequency correlations (see Fig. 1D-G for a visualization, and **Supplementary Figure 1** for spectrograms of all stimuli). Subband means and variances were modified together, referred to as a *base*. For instance, the subband means came from “Bubbling_water.wav” (which is also referred to as the frequency marginal), the subband variances came from “Drumroll.wav” and the cross-frequency correlation values in the high correlation case were taken from “Jogging_on_gravel.wav”. The complete list of source sounds to constrain different properties of the textures is provided in **Table 1** and are provided in the code and data repository (due to size restrictions for the supplementary audio). In total, we created 16 different textures, which comprised all combinations of the constrained properties, i.e. 4 bases (denoted *1.1, 1.2*, *2.1, 2.2*), 2 cross-frequency correlations (low/high CFC), and 2 realizations. The bases X.1 and X.2 differed only by random, amplitude-balanced changes in their subband means, which mimicked the differences we used in a previous study (39). This effectively introduced large (1.X vs. 2.X) and small (X.1 vs X.2) differences in the marginals. As in the study of (18), the CFCs are constrained as marginal (time-average) values between pairs of frequency channels. The average pairwise subband correlation in the low CFC case was 0.2, and in the high CFC case 0.8, inherited from the correlation structure in their respective source sounds.

**Table 1.** List of source sounds for statistical values imposed on white noise.

| Texture Label | Source sound |
| --- | --- |
| Base 1.1/1.2 | Subband means: Bubbling_water.wav<br>Subband variances: Drumroll.wav |
| Base 2.1/2.2 | Subband means: Shaking_coins.wav<br>Subband variances: Crowd_noise.wav |
| Low CFC | Cross-frequency correlations: Fast_running_river.wav |
| High CFC | Cross-frequency correlations: Jogging_on_gravel.wav |

We thus generated a well-controlled set of acoustic textures, whose statistical properties were constrained to be naturalistic based on the statistics of real sounds, however, for which all combinations existed. In particular, this allows us to consider the changes in cross-frequency correlations to be rather general, because it was applied over 4 different combinations of subband means and variances with 2 realizations for each. Importantly, the hearing range of C57Bl/6 mice at this age range is about 2-40 kHz (110,111), i.e. substantially higher than in humans. In order to avoid using all the stimulus energy in low frequencies that are inaudible to the mouse, we applied the statistical properties along the frequency axis in the range 4-64kHz, using the same number of frequency channels (30) as in the statistical analysis of the original sound. The combination of the statistical properties from different sources, and the shift in frequency had the useful effect that the textures were only naturalistic, as opposed to naturally occurring and thus overall novel for the mice and required them to estimate their statistics in the experiment.

Texture synthesis from these statistics was carried out using the above-mentioned toolbox from (18). Briefly, this process entailed modifying a randomly generated Gaussian white noise sample to eventually match a set of prescribed statistics, using a gradient descent based line search. This process was iterative and was terminated when the resulting sound was sufficiently matched for the specified statistic. More details on how textures are synthesized from specified statistical values can be found in (18).

Mouse vocalizations were modeled as inverted ’u’ shaped vocalizations, also referred to as chevron-type by some authors, which are one of the most common types of mouse vocalizations (33,34,112). These vocalizations started with an ascending frequency line, which reached a peak and then decayed more slowly and to a lower frequency than the initial frequency, as is typical for these vocalizations in mice (although a lot of variability exists). Vocalization base frequencies were varied pseudorandomly between 5 frequency locations ([2,4,8,16,32] kHz, see **Supplementary Figure 5C** for marginal spectra of the vocalizations) to ensure animals listened to the presented sounds broadly instead of focusing on a particular frequency range. The sound level of the vocalizations was adapted to a fixed dB difference relative to the subband marginal for each texture, to guarantee a constant degree of difficulty independent of the combination of texture and vocalization frequency. Each vocalization had a frequency range of 0.2 octaves. The parametric design of our background textures (see above) guaranteed that the subband marginal properties (mean and variance) were matched between low and high CFC sounds and therefore any changes in vocalization detection performance had to rely on differences in CFC. In particular, the matched levels between low and high CFC stimuli also did not provide an advantage in relation to the vocalizations, which would randomly be masked strongly or little in high CFC, while more consistently over time in the low CFC case, but to same degrees on average.

To understand the effect of exposure time on the detection of vocalizations, they were presented randomly at 9 linearly spaced onset times from 0 to 6 s. Vocalizations were delivered as a sequence of 10 repetitions at each trial, with an individual vocalization lasting 0.1 s with pauses between adjacent vocalizations lasting 0.1 s (in total, 1.9 s vocalization period). The texture sound continued after the last vocalization for another second. Recent research has demonstrated that mice show a preference for such regularly spaced sequences, while caring less about the precise shape of the vocalizations (113).

Synthesized sounds were converted to an analog output voltage (USB-6351, National Instruments), amplified (A-9030, Onkyo) and delivered at a sampling rate of 250 kHz using a high-fidelity speaker (T250D, Fostex) that had been calibrated before the experiment to output the same sound intensity within 5 dB within the range of 2-100 kHz using an inverse impulse response measured via an ultrasonic microphone (CM16/CMPA, Avisoft).

### Animal training

Experiments were carried out on head-fixed animals inside a sound-proofed booth. The booth was constructed from 3 cm composite wood and the inside walls, floor, and ceiling were covered with acoustic foam (thickness: 5 cm, black surface Basotect Plan50, BASF). The acoustic foam shields against external noises above ∼1 kHz with a sound absorption coefficient >0.95 (defined as the ratio between absorbed and incident sound intensity), which corresponds to >26 dB of shielding in addition to the shielding provided by the booth itself. Sound delivery, data acquisition, and control of the peripherals were performed by custom written software in MATLAB via a data acquisition card (USB-6351, National Instruments).

The purpose of the detection task we used was to teach animals to respond to vocalizations by licking the water source. Licks were automatically registered when the animal’s tongue broke an infrared beam connecting the infrared (IR) light source to the detector located opposite it. The training procedure started with 1-3 days of accommodation to the setup. During this period, animals got used to being head-fixed and rewarded from the water delivery system. At the initial stage of the experimental training, sounds were generated using a set of parameters that made the task easy for the animal. This primarily involved using shorter vocalization onset times, longer vocalization epochs, and higher signal-to-noise ratio (SNR) in the generation of sounds. The difficulty of the task was gradually increased to the level where the vocalizations became non-trivial to detect and potential alternative strategies for carrying out the task did not yield successful outcomes for the animal. To motivate animals to engage in the behavioral task, the total water amount available to any given animal was limited to 1.5 mL/day by an automatic water delivery system that tracked the amount of water delivered in the cage and during the experiment.

Licking before vocalization onset was considered a *false alarm* (FA) and resulted in a penalty waiting period (7 seconds). Licking during the vocalization period was considered a *hit* and was rewarded with 5-7 µL water by opening a solenoid valve (#003-0120-900, Parker). Licking after the last vocalization but before the end of the trial were considered a *miss* (see Fig. 3 for an analysis of these licks). The choice behavior of the animal was tracked over the whole training period to verify that the animal was indeed responding to the vocalizations, rather than using some other task-irrelevant cues. The animals were considered ready for electrode implantation once they displayed a consistently high correlation between first lick time and vocalization onset, showing that their responses were often time-locked to vocalization onset. During the neural recording sessions, the active condition was always run before the passive condition to avoid frustrating the thirsty animals and leading to behavioral extinction.

## Data analysis

### Analysis of neural activity

Neural recordings were bandpass-filtered (0.1-7000 Hz, Butterworth filter, 4th order) and spike-sorted automatically under human control using a custom-written spike-sorter (AutoSort) in Matlab. Sorting was performed per prong of the recording array to better disambiguate neurons visible on multiple electrodes. AutoSort applies a standard sequence of steps, i.e. detection of spikes based on a negative size threshold per recording channel, collection of waveforms from all electrodes on a prong, dimensionality reduction (PCA) to 6 dimensions, followed by clustering (hierarchical clustering with Ward distance) and semiautomatic fusion of clusters based on intra- and between cluster distances. Subsequent analysis only worked on spike timing.

Peristimulus-time histograms (PSTHs) were constructed with different temporal alignments, specifically to texture sound onset, vocalization onset and lick times (see Fig. 3 and 5). Different PSTHs were also constructed for passive and active conditions, as well as different trial outcomes. PSTHs represent the average, normalized firing rate across a set of neurons, i.e. to account for different baseline firing rates, each neuron contributed to these PSTHs normalized by its spontaneous rate, collected across all conditions. In particular this retains differences between active and passive conditions in spontaneous and driven activity. Normalization per alignment can also be informative to highlight relative changes, which we provide in **Supplementary Figure 2**. In the analysis for Fig. 5C, we focussed on vocalization responsive cells, which were identified via a Wilcoxon signed ranks test (p<0.05) between their onset and sustained response.

### Unit classification

To classify units we ran short optical stimulation-only recordings every day prior to sessions involving sound delivery. We used light-evoked response within the 30 ms of light onset and classified units as PV/SST interneurons when the light-evoked response of the unit was significantly higher than the pre-light period. We used a relatively long time window (30 ms) for classification to avoid misclassifying units due to light stimulation artifacts. However, shorter classification windows produced comparable results (**Supplementary Figure 6**). Qualitatively, our classification results were in agreement with previous studies that also characterized spike parameters such as spike width and repolarization time (114–116). We observed that while PV interneurons had features commonly described as fast or narrow spiking (FS), not all cells with narrow spikes crossed the threshold mentioned. Therefore we additionally trained a binary classification model to classify a cell either as FS or not, based on its spike parameters, and analyzed the responses of this group of cells in parallel to optically identified cells.

### Behavioral data analysis

For time-sensitive d’ analysis we combined response times and trial outcomes from all trials for a given animal. Then with 0.75 s time steps, and 1.9 s window size we estimated the d’ for each time point using the approximation d’(t)=Z(HR(t)) - Z(FAR(t)), where Z(p) is the inverse of the Gaussian cumulative distribution function (CDF), HR(t) corresponds to the number of hit and correct rejection trials within the time window, whereas FAR(t) corresponds to the number of false alarms. For all other behavioral analyses hit rate was estimated either at the level of session or animal was used. The hit rate for a given condition was simply the ratio of hit trials to total trials within a given condition. For tracking the training process, we shuffled the reaction times for a given session and estimated the mean shuffled hit rate using 100 randomizations. The successful training session was defined as the one with a hit rate > *μ_shuffled_ +* 2*σ_shuffled_*.

### Stimulus reconstruction

To quantify the information in the neural population we performed stimulus reconstruction from population response under passive recording conditions using the "flat prior reconstruction" method described earlier (46). In contrast to the "optimal prior reconstruction" method from the same study the flat prior method removes the correlations from the input in the STRF estimation step and therefore does not have an inherent advantage for sounds with higher cross-frequency correlations, which would have confounded out comparison of reconstructions across the different levels of CFCs. Specifically, first, a subset of units (395 out of 1353) was selected based on the signal-to-noise ratio (SNR) of their STRF. Linear reconstruction kernels were computed using training data that encompassed the background only section of the balanced set of stimuli. Kernels were estimated this way 20 times using different sets of stimuli, randomly selected from the training data and the final reconstruction kernels used in the analysis was the average across these randomizations. Reconstruction kernels were then projected onto the neural responses from the test dataset in order to obtain reconstructed spectrograms. The quality of reconstruction was quantified using point-wise correlation between the original and reconstructed spectrograms.

### Vocalization decoding with support vector machine (SVM) analysis

A binary SVM classifier was used for classifying a given population vector as belonging to the vocalization (1) or background epoch (0). The model was trained using a subset of data balanced for correlation, light stimulation, and vocalization base frequencies. Spike counts for each time point and the unit, SC(t,u), from the training dataset, were divided into SC(t_VOC_, u) and SC(t_BKG_, u) parts corresponding to data coming from vocalization and background epochs. The trained model was then 10-fold cross-validated. The model was used for labeling population vectors belonging to the vocalization epoch of the test dataset. Since vocalizations were repeated multiple times during the trial (See Acoustic stimulus section), the probability of being classified as vocalization was averaged across different repetitions to yield a single mean time series of 200 ms length. However, statistical tests (Kruskal-Wallis test) between different classification probabilities were performed on unaveraged, raw probabilities outputted by the SVM classifier.

### Factorization of variance with nested ANOVA

The variance of unit responses into texture class and realization components was performed as described in detail earlier (54). For this analysis, we averaged binned spike counts for each cell using moving to mean with window sizes ranging from 30 ms to 1000 ms. The resulting neural response matrix for each cell had dimensions of n(stimuli)×n(time bins) but the endpoints of the matrices were discarded, yielding shorter matrices when the window size was larger. Spike counts were normalized to the baseline period before the sound onset. To analyze the variance of the cell’s response at a given time point, nested ANOVA was performed. Realizations (2 per class) were excerpts taken from different parts of the same clip. Each clip belonged to a certain texture class (8 in total) defined by its marginal statistics and cross-frequence correlation. The realization variable was nested under the texture class. Following (54) we used the variance explained (VE) by the ANOVA model to understand the effect of window size, time course, and cell type on the analysis. Population VE by each variable was obtained by averaging across cells. To summarize the relationship between the window size and VE, we further averaged the population VE across time points.

### Convergence of Statistical Properties

To check whether low and high CFC stimuli differed in the variability of contained statistics in shorter duration samples, we ran excerpts with durations ranging from 100 ms to 10 s through the estimation routines provided by (18), using the function measure_texture_stats. We used 10 samples per duration and realization. The resulting statistics were collected and next the standard deviation across the estimates was computed for the envelope marginals, envelope variance, cross-frequency correlations and modulation power in each frequency band. The results of this analysis are shown in **Supplementary Figure 4**.

## Acknowledgements

We would like to express our gratitude to Amber van der Stam, Dionne Lenferink, and Tom van Iersel for their assistance with animal training. BE also acknowledges valuable discussions at the Kavli Conference on Statistical Learning in the Brain at UCSB, supported by NSF Grant No. PHY-1748958 and the Gordon and Betty Moore Foundation Grant No. 2919.02 to the Kavli Institute for Theoretical Physics (KITP). The present work is based on Chapter 2 of the first author’s PhD thesis *Processing of Statistically Defined Sounds in the Auditory Cortex* (117).

## Author contributions

Conceptualization: AA, KP, BE

Methodology: AA, KP, PvH, BE

Investigation: AA, KP, PvH, BE

Software: AA, KP, PvH, BE

Data curation: AA, KP, PvH, BE

Validation: AA, KP, PvH, BE

Supervision: AA,BE

Resources: AA, BE

Formal analysis: AA

Visualization: AA, BE

Writing—original draft: AA, BE

Writing—review & editing: AA, KP, PvH, BE

Project administration: BE

Funding acquisition: BE

## Data and Materials Availability

All data needed to evaluate the conclusions in the paper are present in the paper and/or the Supplementary Materials. The code used for generating the figures in this manuscript are available in the Zenodo Repository at https://doi.org/10.5281/zenodo.13779387

## Supplementary Information

**Supplementary Figure 1.**
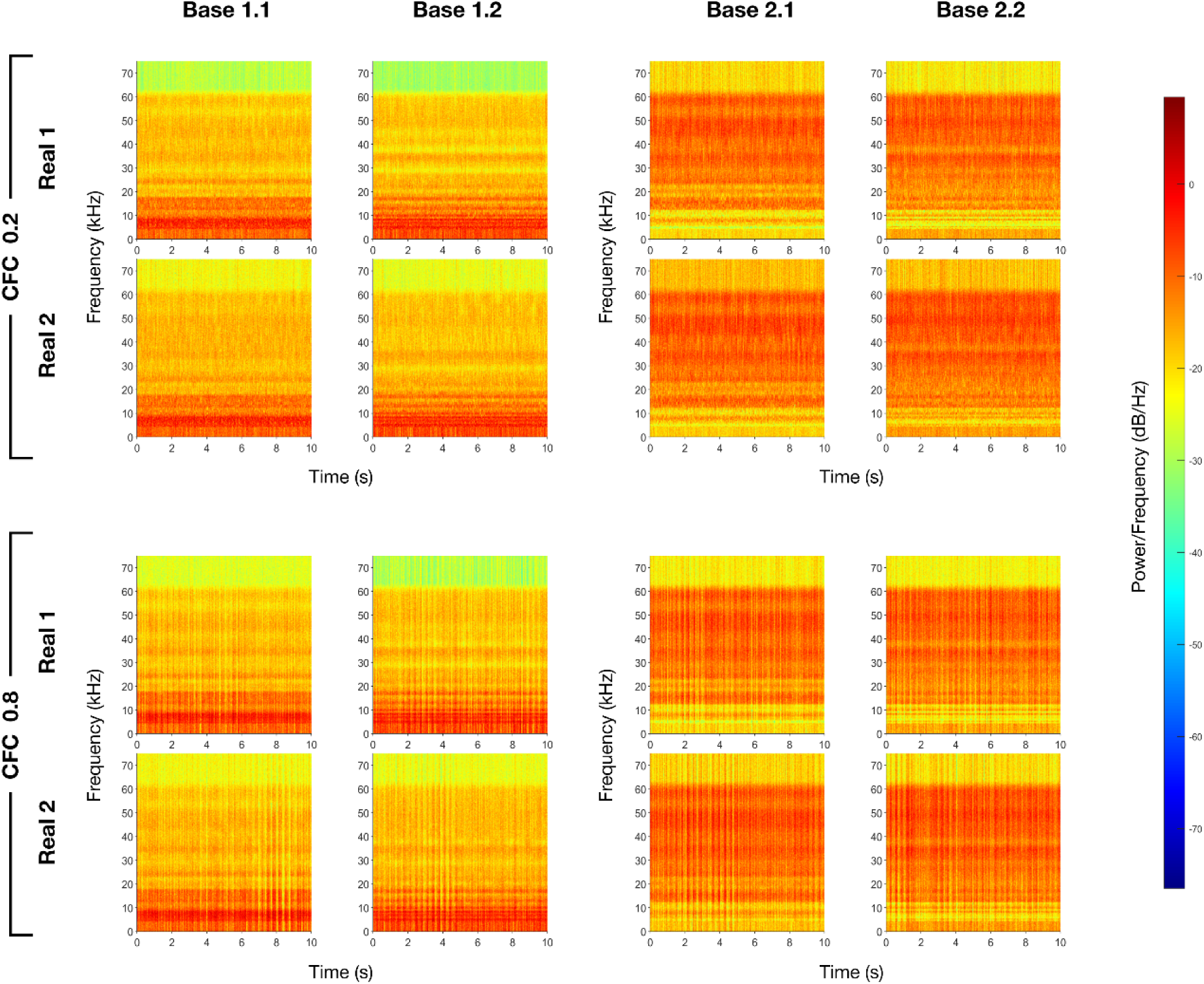
Spectrograms of background textures used in the experiment. In total, 16 distinct background textures were generated and used in the experiment. The textures were created using two different base frequency configurations (Base 1 and Base 2, each with two variants: 1.1, 1.2, 2.1, and 2.2, columns), defined by their envelope means and variances. Bases 1 and 2 were chosen to cover low and high frequencies, in order to activate a large set of neurons. The textures were further differentiated by their cross-frequency correlation (CFC, top/bottom rows), with either high (0.8) or low (0.2) average correlation across frequency bands. For each combination of CFC and base configuration, two independent realizations (Real 1 and Real 2, rows within CFCs) were generated.

**Supplementary Figure 2.**
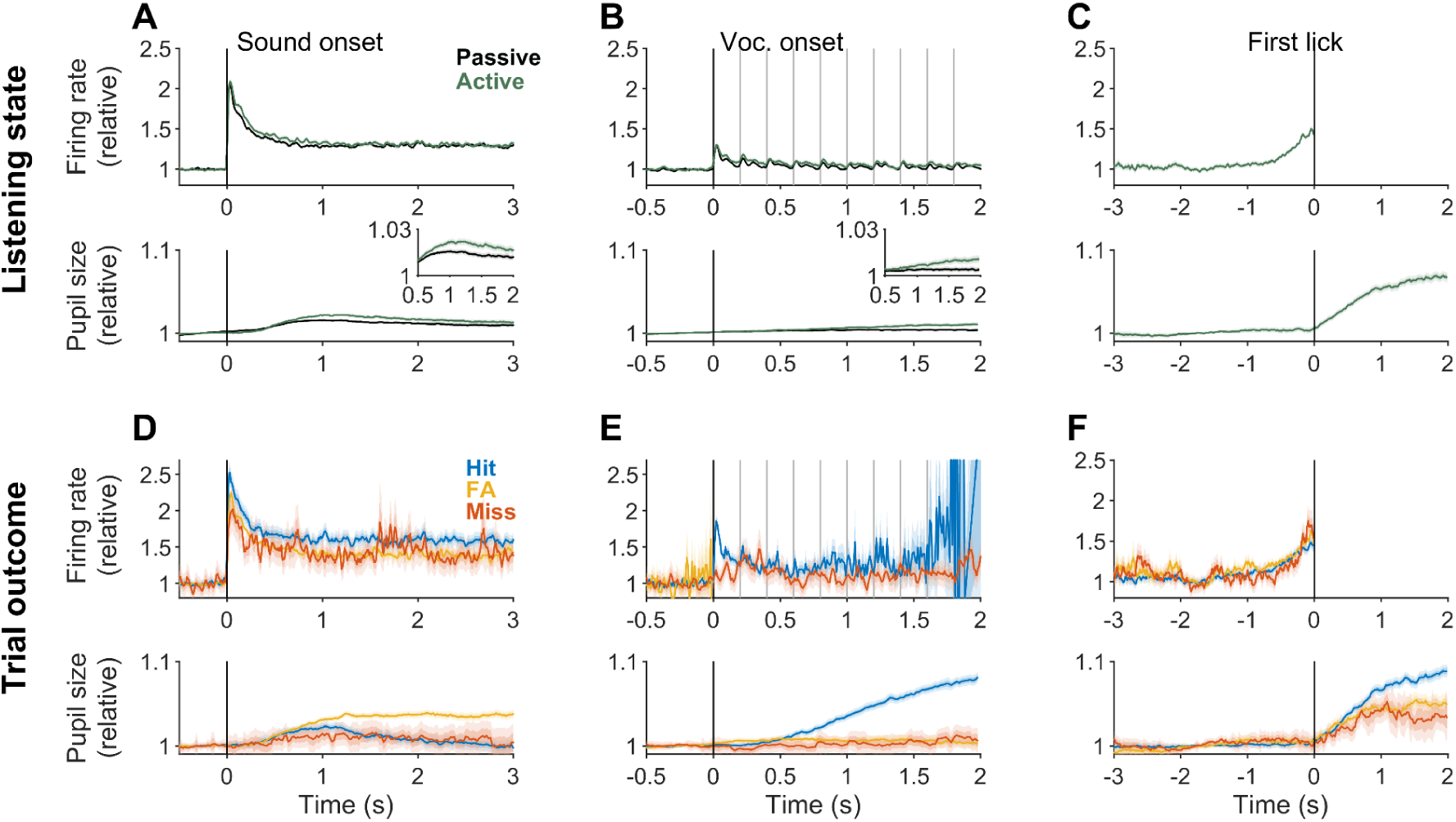
Unit responses normalized by experimental condition. Similar to Figure 3 in the main text, but here the response was normalized separately for each condition. For sound and vocalization onset-aligned responses, the 0.5 s time window prior to the respective event was used to normalize the data. For first-lick aligned responses, the time window from -2 to -1.5 s prior to the lick was used to normalize the data.

**Supplementary Figure 3.**
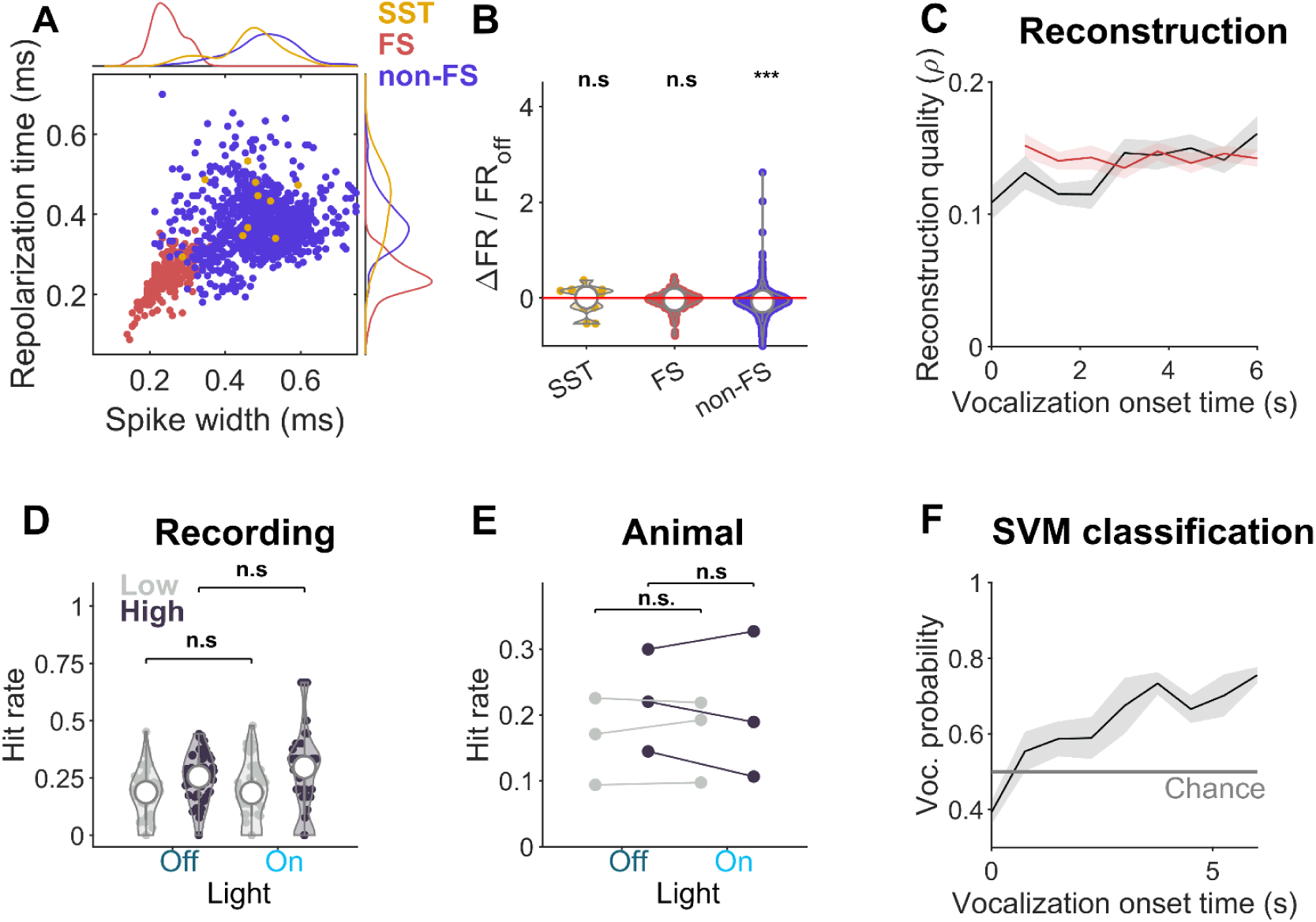
Manipulation of SST cells does not affect behavioral performance. **A** Unlike PV cells shown in Figure 6, SST cells (n = 12) were found to overlap primarily with non-FS cells. **B** Activation of SST cells does not affect the sound-evoked activity of SST (median = 1.67%, Wilcoxon signed rank test, p = 0.791) and FS cells (median = 1.35%, Wilcoxon signed rank test, p = 0.752), but significantly reduces that of non-FS cells (median = -3.82%, Wilcoxon signed rank test, p < 0.001). **C** Reconstruction quality of vocalization (light gray) and background (red) epoch is quantified for stimuli with different vocalization onset times. The thick black line represents average reconstruction quality across sounds with identical vocalization onset times. **D-E** Activation of SST cells does not affect the behavioral performance for low or high CFC backgrounds, neither at recording (n = 60 recordings; Wilcoxon signed rank test, p = 0.697 for low CFC; p = 0.416 for high CFC sounds) nor at animal levels (n = 3; Wilcoxon signed rank test, p = 0.750 for low CFC; p = 0.500 for high CFC). **F** The probability of the population vector being classified as coming from the vocalization epoch gradually increased in SST dataset as well. Solid gray line represents chance-level performance for the SVM classifier.

**Supplementary Figure 4.**
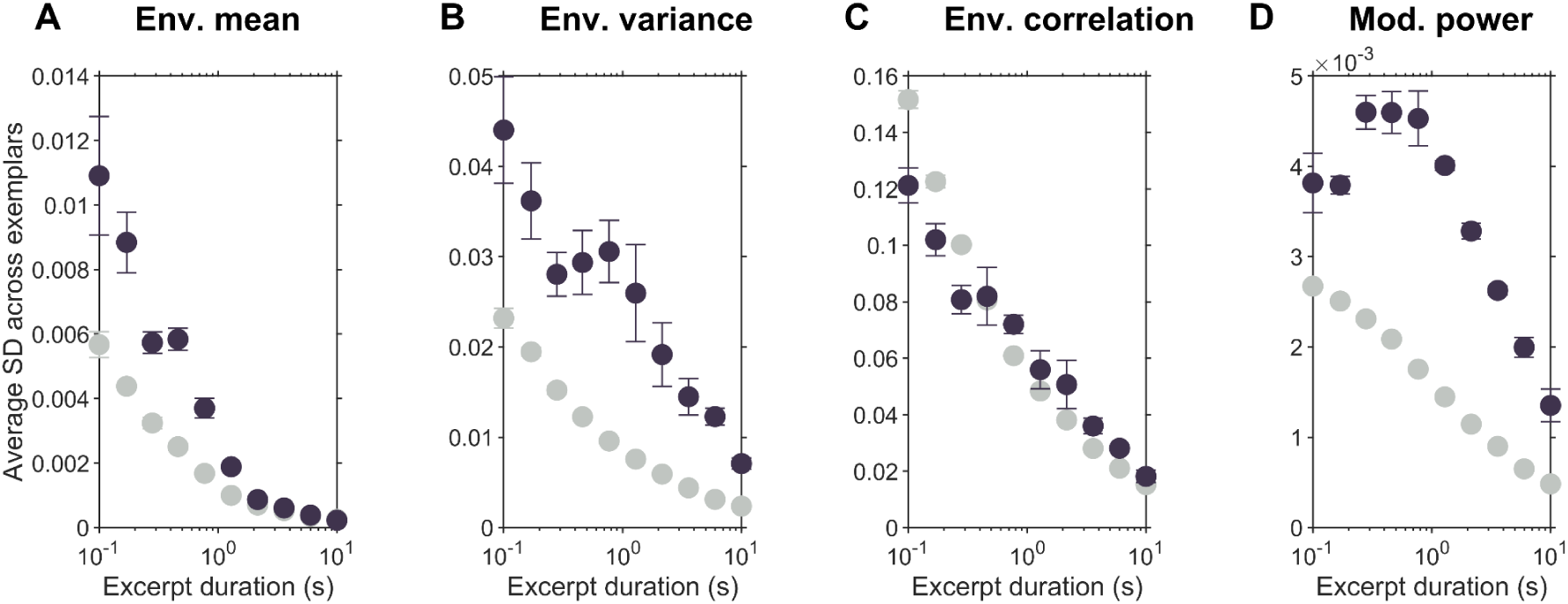
Statistics time-courses for low and high CFC textures. We used the toolset of (*18*) to analyze the variability in statistical properties estimated from different lengths of data, ranging from 100 ms to 10 s, i.e. covering the range of durations used in our experiment. Importantly, the values below are the S.D. across 10 samples, i.e. are not the value for the statistic itself, mirroring the analysis in Fig. 1C, (*25*) Error bars show SEM across textures. **A** The marginals of the envelopes per frequency channel converged after ∼1 s, with low CFC textures exhibiting a faster decay than high CFC textures, which is consistent with the greater variability across time in the latter. **B** The variance of the envelope per frequency channel converged more slowly than the marginals, with CFC textures converging much more slowly and exhibiting more residual variability. **C** The correlation across frequency channels exhibited also slower convergence of estimation accuracy with duration, which however, did not markedly differ between low and high CFCs. **D** The modulation power in the envelopes surprisingly exhibited a non-monotonic decay, for high CFC stimuli only, suggesting that new time-scales of modulation came ’into sight’, when the duration increased to around 500 ms, before then converging. Low CFC textures instead showed a slow decay that was comparable in speed to envelope variance and correlation.

**Supplementary Figure 5.**
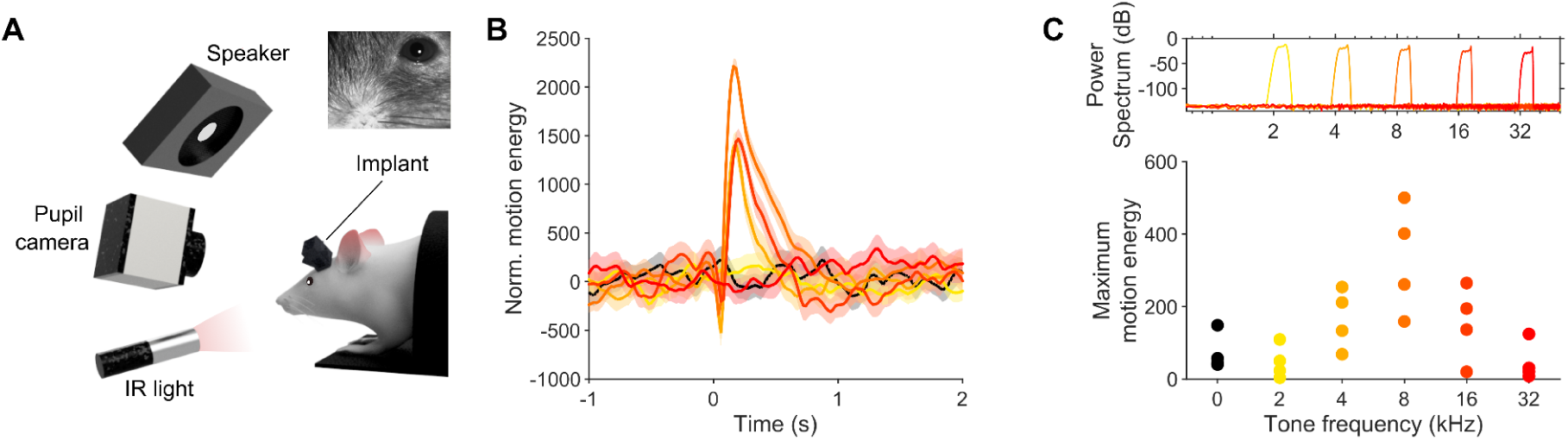
Quantifying hearing in untrained animals through facial motion analysis. **A** Untrained animals (n = 4) were exposed to a randomized sequence of pure tones (t = 2 s), while the animal’s face was recorded (inset). **B** Recorded video data was subjected to dimensionality reduction via singular value decomposition (SVD). The motion energy of the first component, normalized to different tone frequencies (from yellow to red), is shown here, aligned to the tone onset for an example session. **C** Top: Envelope mean of vocalization sounds showing power across different frequencies (2-32 kHz) for vocalizations used in the task. Bottom: The maximum response of the first component to different frequencies demonstrates that the animal’s response to certain frequencies is reduced to the level of the control condition of silence (black dots), indicating that frequencies on the edges of hearing ability (2 and 32 kHz) are no longer heard as well as frequencies between 4-16 kHz.

**Supplementary Figure 6.**
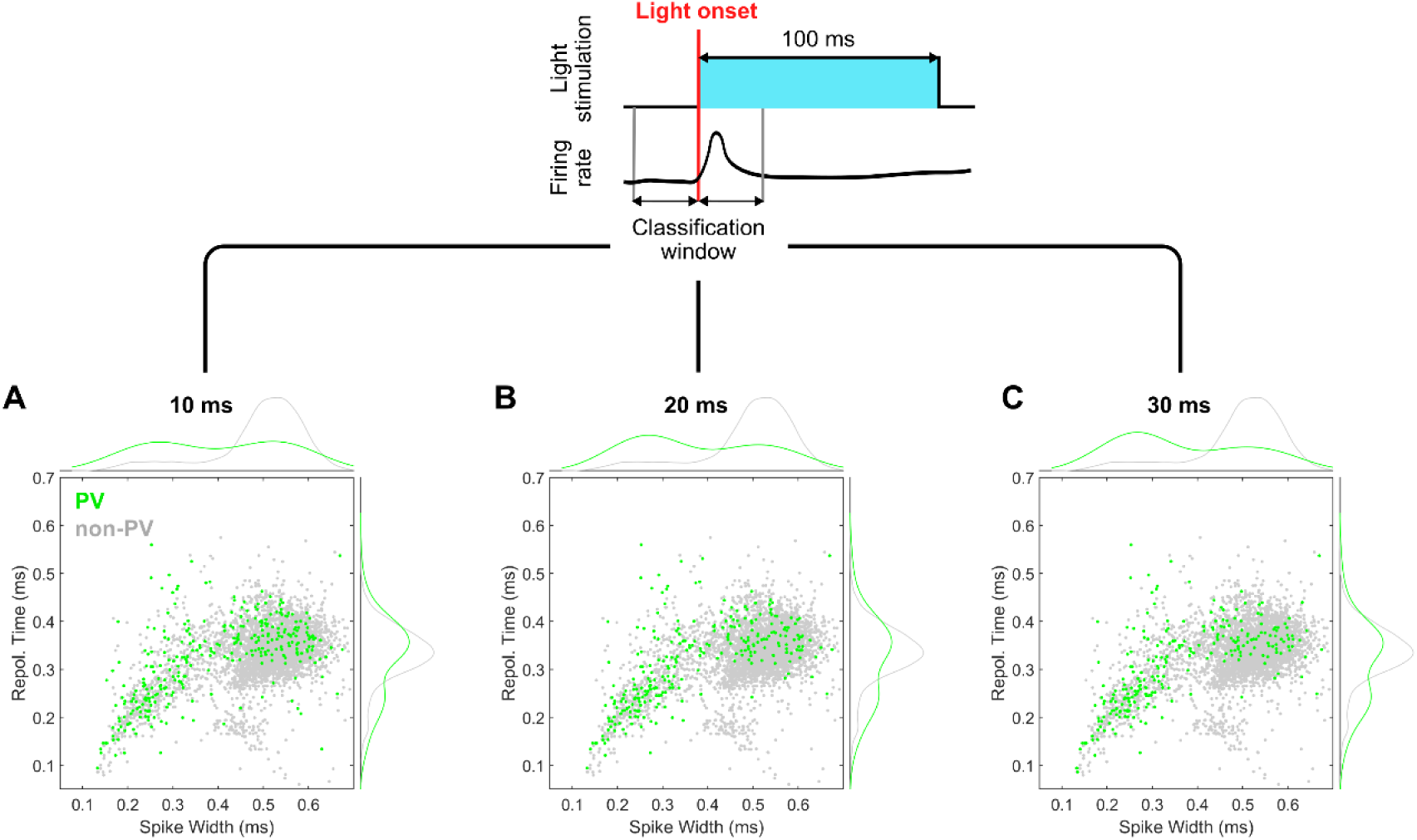
Classification of light-sensitive neurons using different temporal windows. Individual neurons were classified as PV (green) or non-PV (gray) based on their responses to light stimulation. Classification was performed by statistically comparing firing rates before and after light onset during repeated stimulation trials. Panels A, B, and C show the distribution of classified neurons when using classification windows of 10 ms, 20 ms, and 30 ms respectively around the light onset. While the number of identified PV neurons decreases with longer classification windows (A: n=336; B: n=293; C: n=264), the distribution of spike parameters (repolarization time vs. spike width) remains quite similar across all temporal windows. 78% of cells classified as PV cells with a 30 ms classification window are also classified as such using shorter window lengths (in contrast to 61% and 71% classified using 10 and 20 ms, respectively). Density plots along the axes show the distribution of each parameter for both cell types.

